# Multimodal modeling of neural network activity: computing LFP, ECoG, EEG and MEG signals with LFPy2.0

**DOI:** 10.1101/281717

**Authors:** Espen Hagen, Solveig Næss, Torbjørn V. Ness, Gaute T. Einevoll

## Abstract

Recordings of extracellular electrical, and later also magnetic, brain signals have been the dominant technique for measuring brain activity for decades. The interpretation of such signals is however nontrivial, as the measured signals result from both local and distant neuronal activity. In volume-conductor theory the extracellular potentials can be calculated from a distance-weighted sum of contributions from transmembrane currents of neurons. Given the same transmembrane currents, the contributions to the magnetic field recorded both inside and outside the brain can also be computed. This allows for the development of computational tools implementing forward models grounded in the biophysics underlying electrical and magnetic measurement modalities.

LFPy (LFPy.readthedocs.io) incorporated a well-established scheme for predicting extracellular potentials of individual neurons with arbitrary levels of biological detail. It relies on NEURON (neuron.yale.edu) to compute transmembrane currents of multicompartment neurons which is then used in combination with an electrostatic forward model. Its functionality is now extended to allow for modeling of networks of multicompartment neurons with concurrent calculations of extracellular potentials and current dipole moments. The current dipole moments are then, in combination with suitable volume-conductor head models, used to compute non-invasive measures of neuronal activity, like scalp potentials (electroencephalographic recordings; EEG) and magnetic fields outside the head (magnetoencephalographic recordings; MEG). One such built-in head model is the four-sphere head model incorporating the different electric conductivities of brain, cerebrospinal fluid, skull and scalp.

We demonstrate the new functionality of the software by constructing a network of biophysically detailed multicompartment neuron models from the Neocortical Microcircuit Collaboration (NMC) Portal (bbp.epfl.ch/nmc-portal) with corresponding statistics of connections and synapses, and compute *in vivo*-like extracellular potentials (local field potentials, LFP; electrocorticographical signals, ECoG) and corresponding current dipole moments. From the current dipole moments we estimate corresponding EEG and MEG signals using the four-sphere head model. We also show strong scaling performance of LFPy with different numbers of message-passing interface (MPI) processes, and for different network sizes with different density of connections.

The open-source software LFPy is equally suitable for execution on laptops and in parallel on high-performance computing (HPC) facilities and is publicly available on GitHub.com.

## 1. Introduction

Ever since the 1950s, electrical recordings with sharp electrodes have been the most important method for studying *in vivo* activity in neurons and neural networks [Li and Jasper, 1953]. In the last couple of decades, however, a host of new measurement methods has been developed and refined. One key development is the new generation of multicontact electrodes allowing for high-density electrical recordings across cortical laminae and areas, and the accompanying resurgence of interest in the low-frequency part of the extracellular signal, the ‘local field potential’ (LFP) [Buzsáki, 2004; Buzsáki et al., 2012; Einevoll et al., 2013]. The LFP is a population measure reflecting how dendrites integrate synaptic inputs, insight that cannot be obtained from measurement of spikes from a handful of neurons [Einevoll et al., 2013]. Many new optical techniques for probing cortical activity have also been developed. Of particular interest is *two-photon calcium imaging*, which can measure the action potentials of individual neurons deep into cortical tissue [Helmchen and Denk, 2005], and *voltage-sensitive dye imaging (VSDI*), which measures the average membrane potential across dendrites close to the cortical surface [Grinvald and Hildesheim, 2004]. These add to the more established systems-level methods such as *electroencephalography* (*EEG*, Nunez and Srinivasan [2006]), which measures electrical potentials at the scalp, and *magnetoencephalography* (*MEG*, Hämäläinen et al. [1993]) which measures the magnetic field outside the head.

A standard way of analyzing such neurophysiological data has been to look for correlations between measurements and how the subject is stimulated or behaves. For example, most of what we have learned about neural representation of visual information in visual cortex has come from receptive-field studies where the correlation between measured spikes and presented visual stimuli is mapped out [Hubel and Wiesel, 1959]. The same approach has been used to map out the receptive fields for other sensory modalities (sound, touch, etc.), objects and celebrities [Quiroga et al., 2005], or the spatial location of the animal [O’Keefe and Dostrovsky, 1971; Hafting et al., 2005].

This purely statistical approach has limitations, however. For one, it only provides estimates for the neural representation and gives no direct insight into the circuit mechanisms giving rise to these representations. Secondly, the receptive field is inherently a *linear* measure of activity [Dayan and Abbott, 2001] and cannot in general capture non-linear network dynamics. The receptive field in primary visual cortex depends, for example, strongly on stimulation of the surrounding regions of visual space, an inherently non-linear effect [Blakemore and Tobin, 1972]. For other cortical measurements, such as the LFP or VSDI, a statistical analysis is further complicated by the fact that the signals reflect activity in neuron populations rather than individual neurons [Petersen et al., 2003; Einevoll et al., 2013]. This makes commonly-used statistical signal measures such as power spectra, correlation, coherence, and functional connectivity difficult to interpret in terms of activity in neurons and networks [Einevoll et al., 2013].

An alternative approach to a purely statistical analysis is, following in the tradition of physics, to formulate candidate hypotheses precisely in mathematics and then compute what each hypothesis would predict for the different types of measurements. Until now candidate cortical network models have typically only predicted spiking activity, thus preventing a proper comparison with measurements other than single-unit and multiunit recordings. To take full advantage of all available experiments, there is a need for biophysics-based forward-modeling tools for predicting other measurement modalities from candidate network models [Brette and Destexhe, 2012], that is, develop software that faithfully models the various types of measurements themselves. To facilitate the forward-modeling of extracellular potentials, both LFPs and spikes (i.e., either single-unit or multi-unit activity (MUA)), we developed LFPy (LFPy.readthedocs.io, Lindén et al. [2014]), a Python tool using the NEURON simulator [Carnevale and Hines, 2006] and its Python interface [Hines et al., 2009].

LFPy implements a well-established forward-modeling scheme where the extracellular potential is computed in a two-step process [Holt and Koch, 1999; Lindén et al., 2014]: First, the transmembrane currents of multicompartment neuron models are computed using NEURON. Second, the extracellular potential is computed as a weighted sum over contributions from the transmembrane currents from each compartment with weights prescribed by volume-conductor theory for an infinite volume conductor. In LFPy these functions are provided by a set of Python classes that can be instantiated to represent the cell, synapses, stimulation devices and extracellular electric measurement devices. By now this forward-model method has been used in a number of studies, for example to model extracellular spike waveforms [Holt and Koch, 1999; Gold et al., 2006, 2007; Pettersen and Einevoll, 2008; Pettersen et al., 2008; Franke et al., 2010; Schomburg et al., 2012; Thorbergsson et al., 2012; Reimann et al., 2013; Ness et al., 2015; Hagen et al., 2015; Miceli et al., 2017; Cserpán et al., 2017], LFP signals [Pettersen et al., 2008; Lindén et al., 2010, 2011; Gratiy et al., 2011; Makarova et al., 2011; Schomburg et al., 2012; Łęski et al., 2013; Reimann et al., 2013; Martín-Vázquez et al., 2013, 2015; Gła˛bska et al., 2014; Mazzoni et al., 2015; Tomsett et al., 2015; Sinha and Narayanan, 2015; Taxidis et al., 2015; Hagen et al., 2016; Gła˛bska et al., 2016; Ness et al., 2016; Hagen et al., 2017] and recently axonal LFP contributions [McColgan et al., 2017]. Some of these used LFPy to predict extracellular potentials [Łęski et al., 2013; Lindén et al., 2014; Hagen et al., 2015; Tomsett et al., 2015; Mazzoni et al., 2015; Ness et al., 2015, 2016; Hagen et al., 2016, 2017; Miceli et al., 2017], while in Heiberg et al. [2016] LFPy was used to construct a small-world LGN network without predictions of extracellular potentials. Further, in Uhlirova et al. [2016] LFPy was used to compute neuronal membrane potentials.

Here we present a substantially extended version of LFPy, termed LFPy2.0, including several new features, that is, support for (i) simulations of networks of multicompartmental neuron models, (ii) computation of LFP/MUA with anisotropic electrical conductivity, (iii) computation of LFP/MUA in the presence of step-wise varying electrical conductivity (such as at the interface between cortical gray matter and white matter), (iv) computation of ECoG signals (i.e., electrical potentials recorded at the cortical surface), (v) computation of EEG signals, and (vi) computation of MEG signals, see illustration in Fig. 1. To illustrate the computation of these measures by LFPy2.0 we show in Fig. 2 the LFP, EEG and MEG signals generated by a single synaptic input onto a single pyramidal neuron. As both electric and magnetic signals sum linearly, the recorded signals in real applications will stem from the sum of a large number of such contributions.

**Figure 1:**
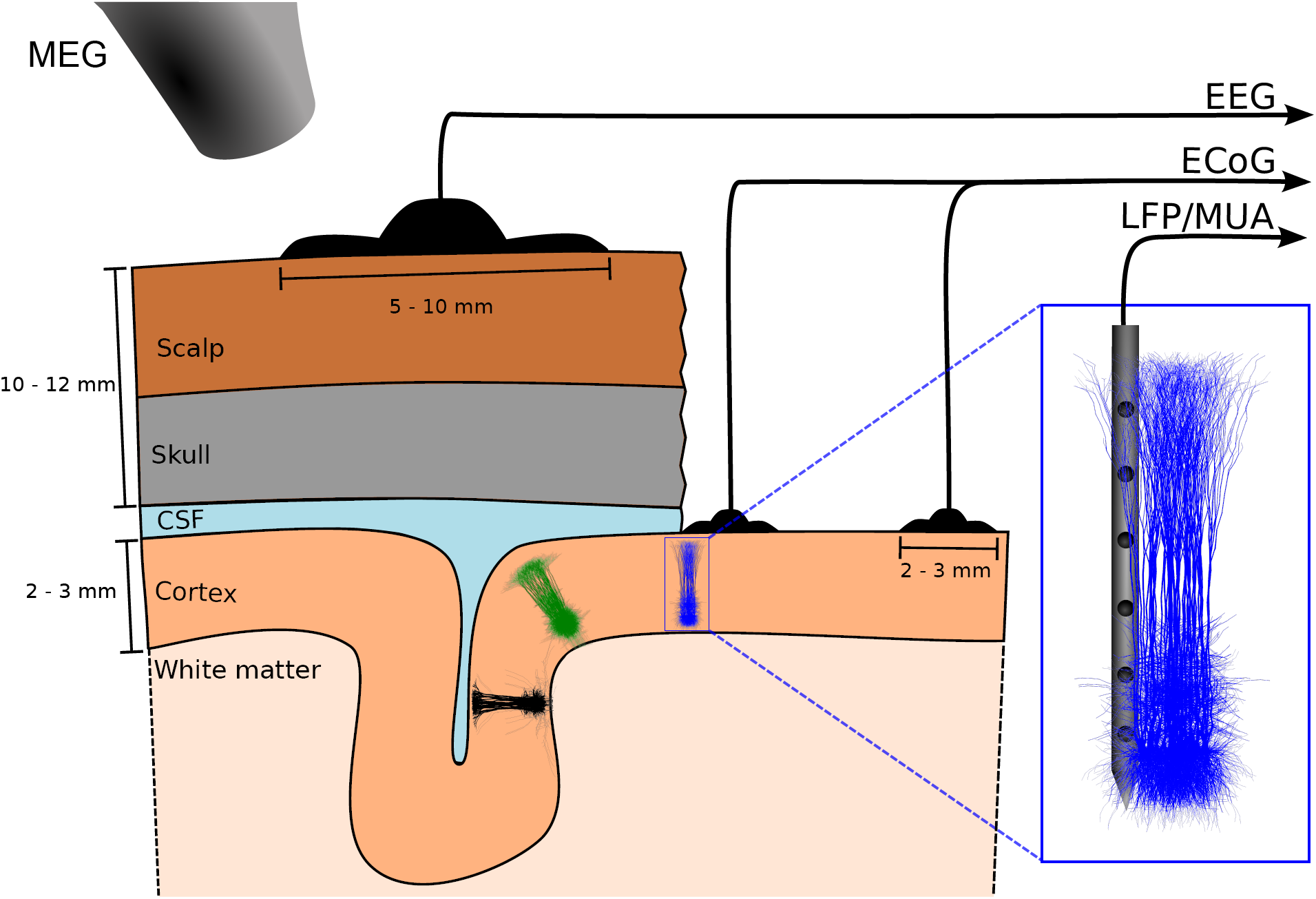
Illustration of measurement signals computed by LFPy2.0. The figure illustrates the EEG, ECoG, LFP/MUA (linear multi-electrode) and MEG recordings of electrical and magnetic signals stemming from populations of cortical neurons. Here three separate cortical populations are depicted. EEG electrodes are placed on the scalp, ECoG electrodes on the cortical surface, while the LFP and MUA both are recorded by electrodes placed inside cortex. In MEG the tiny magnetic fields stemming from brain activity is measured by SQUIDs placed outside the head. The MUA signal, that is, the high-frequency part of the recorded extracellular potential inside cortex, measures spikes from neurons in the immediate vicinity of the electrode contact, typically less than 100 *µ*m away [Buzsáki, 2004; Pettersen and Einevoll, 2008; Pettersen et al., 2008]. The ‘mesoscopic’ LFP and ECoG signals will typically contain information from neurons within a few hundred micrometers or millimetres from the recording contact [Einevoll et al., 2013], while the ‘macroscopic’ EEG and MEG signals will have contributions from cortical populations even further away [Nunez and Srinivasan, 2006; Hämäläinen et al., 1993].

**Figure 2:**
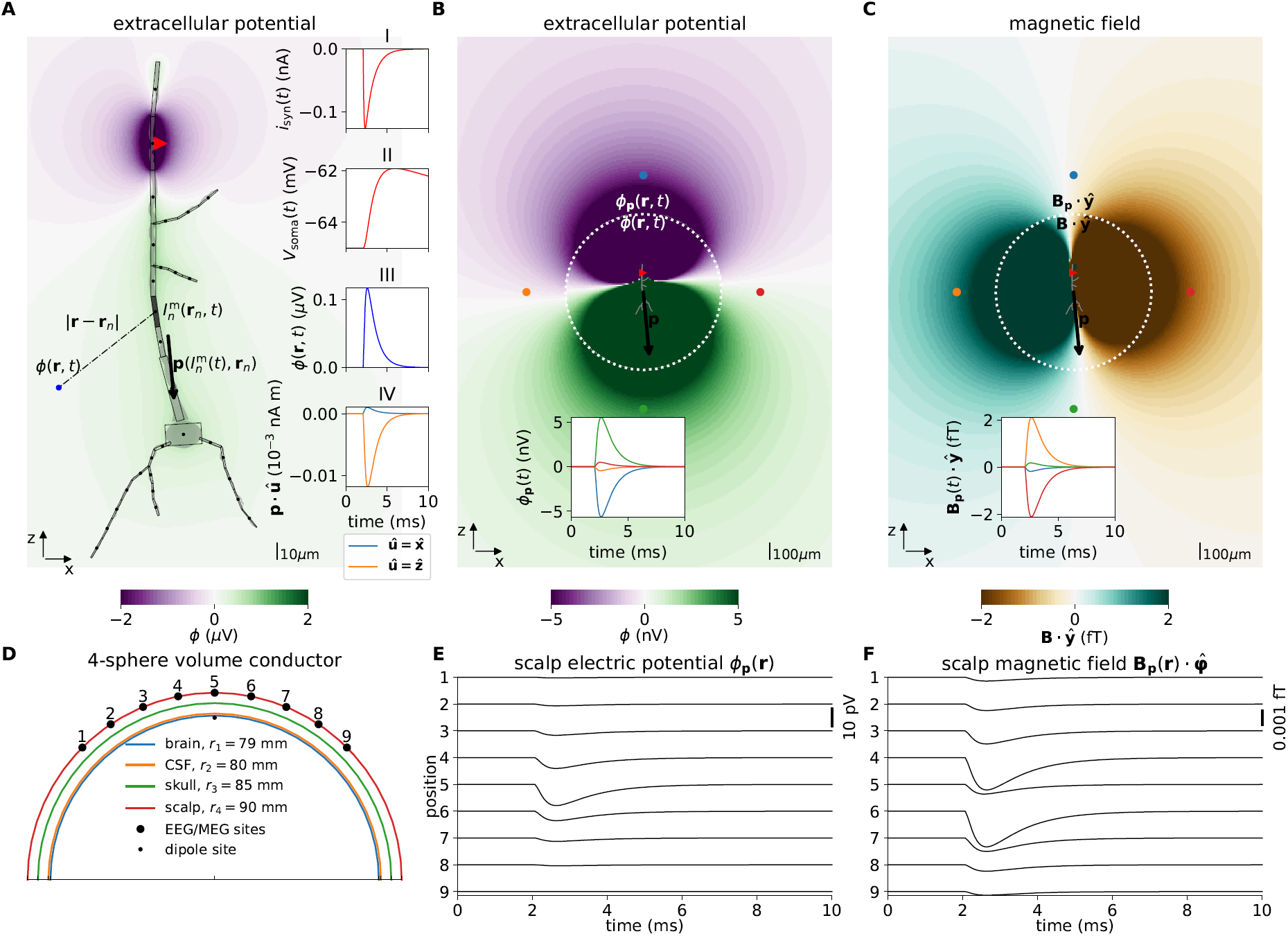
Illustrations of forward model, dipole approximation, EEG and MEG model. **A** Illustration of forward-modeling scheme for extracellular potentials from multicompartment neuron models. The gray shape illustrates a 3D-reconstructed neuron morphology and the equivalent discretized multicompartment model. A single synaptic input current *i*^syn^(*t*) (red triangle, inset axes I) results in a deflection of the membrane voltage throughout the morphology, including at the soma (*V*_soma_(*t*), inset axes II). LFPy allows for computing extracellular potentials *φ* in arbitrarily chosen extracellular locations **r** (inset axes III) from transmembrane currents 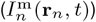, as well as the radial and tangential components of the current dipole moment **p** (black arrow, inset axes IV). Compartments are indexed *n*, **r***n* denote compartment positions. The image plot shows the extracellular potential in the *xz*-plane at the time of the largest synapse current magnitude (*t* = 2.25 ms). **B** Illustration of the extracellular electric potential, calculated both from the current dipole moment and transmembrane currents for the situation in panel A. Within a radius *r <* 500 *µ*m from the ‘center of areas’ (see below) of the morphology the panel shows extracellular potentials *φ*(**r**) predicted using the line-source method, while outside this radius the panel shows extracellular potentials *φ*_**p**_(**r**) predicted from the current dipole moment (**p**, black arrow). Here, an assumption of an infinite, homogeneous (same everywhere) and isotropic (same in all directions) extracellular conductivity was used. The ‘center of areas’ was defined as 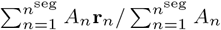 where *A*_*n*_ denotes compartment surface area. The time *t* = 2.25 ms as in panel A. The inset axis shows the potential as function of time in the four corresponding locations (at **R** = 750 *µ*m) surrounding the morphology (colored circular markers). **C** Visualization of magnetic field component **B**_P_ · 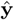 (*y*-component) computed from the current dipole moment, outside a circle of radius *r* = 500 *µ*m (as in panel B). Inside the circle, we computed the same magnetic field component from axial currents inside each compartment. The inset axis shows the *y*-component of the magnetic field as function of time in the four corresponding locations (at **R** = 750 *µ*m) surrounding the morphology (circular markers). **D** Illustration of upper half of the four-sphere head model used for predictions of EEG scalp potentials from electric current dipole moments. Each spherical shell with outer radii *r ϵ{r*_1_, *r*_2_, *r*_3_, *r*_4_} has piecewise homogeneous and isotropic conductivity *σ*eϵ {*σ*_1_, *σ*_2_, *σ*_3_, *σ*_4_}. The EEG/MEG sites numbered 1-9 mark the locations where electric potentials and magnetic fields are computed, each offset by an arc lengsth of *r*_4_*π/*16 in the *xz*-plane. The current dipole position was *θ* = *φ* = 0, *r* = 78 mm (in spherical coordinates). **E** Electric potentials on the outer scalp-layer positions 1-9 in panel D. **F** Tangential component of the magnetic field **B**_P_ · 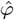 in positions 1-9. (Note that at position 5, the unit vector 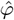 is defined to be directed in the positive *y*-direction.)

The manuscript is organized as follows: In Methods we first review the biophysical forward-modeling scheme used to predict extracellular potentials in different volume-conductor models. Then we describe calculations of current dipole moments and corresponding calculation of EEG and MEG signals. We further describe new LFPy classes and corresponding code examples for set-up of networks, the implementation of an example network using available data and biophysically detailed cell models from the Blue Brain Project’s Neocortical Microcircuit Collaboration (NMC) Portal, and various technical details. In Results we investigate the outcome of our example parallel network simulation and corresponding measurements, and assess parallel performance of LFPy when running on HPC facilities. In Discussion we outline implications of this work and discuss possible future applications and developments of the software.

## 2. Methods

### 2.1. Multicompartment modeling

#### 2.1.1. Calculation of transmembrane currents

The origin of extracellular potentials is mainly transmembrane currents [Buzsáki et al., 2012; Einevoll et al., 2013], even though diffusion of ions in the extracellular space alone also can give rise to such poten-tials [Halnes et al., 2016]. In the presently (and frequently) used forward modeling approach, these trans-membrane currents are obtained from spatially discretized multicompartment neuron models [De Schutter and Van Geit, 2009] which allow for high levels of biophysical and morphological detail. Such models have historically been used to model spatiotemporal variations in the membrane voltages *V* ^m^(*x, t*), where *x* de-notes the position along an unbranched piece of dendritic cable. From this cable theory it also follows that the transmembrane current density, that is, the transmembrane current per unit length of membrane, for any smooth and homogeneous cable section is given by [Koch, 1999]:

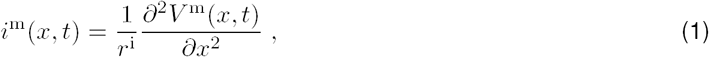

where *r*^i^ represents the axial resistance per unit length along the cable. Assuming a homogeneous current density per unit length *i*^m^ along a single compartment with length Δ*s*, the total transmembrane current *I*^m^ = *i*^m^Δ*s*.

As in the first release of LFPy [Lindén et al., 2014], we rely on the NEURON simulation environment [Carnevale and Hines, 2006] to compute transmembrane currents. As of NEURON v7.4, a faster and direct method of accessing transmembrane currents is provided through its `CVode.use_fast_imem()` method, which we now utilize in an exclusive manner. NEURON’s ‘extracellular’ mechanism is thus no longer used to predict extracellular potentials (cf. Lindén et al. [2014, sec. 5.6]). Note, however, that this mechanism itself is still used when an external extracellular potential is imposed as a boundary condition outside each compartment using the `Cell.insert_v_ext()` class method.

#### 2.1.2. Calculation of axial currents

To compute the magnetic fields stemming from electrical activity in neurons, the axial currents within cells are needed [Hämäläinen et al., 1993]. The axial current for the cable is given by [Koch, 1999]:

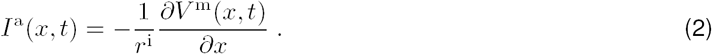

Assuming homogeneous axial current density between the midpoints of two neighboring compartments *n* and *n* + 1 along the cable, one may obtain the axial current from Ohm’s law:

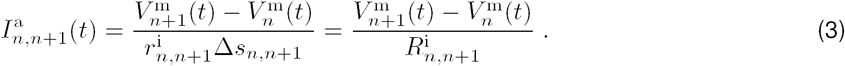

Here, 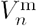 and 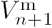 are the compartment midpoint membrane potentials, 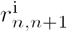 axial resistance per unit length between the two compartments, *Δs_n,n+1_* the distance between compartment midpoints and 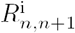 the corresponding axial resistance.

Further, we outline how axial currents from complex reconstructed neuron morphologies are calculated in LFPy2.0, and provide the technical implementation details in Algorithm 1 below. For a more comprehensive explanation, see Næss [2015]. The corresponding implementation is in LFPy2.0 provided by the class method `Cell.get_axial_currents_from_vmem()`.

In NEURON, a *section* is a continuous piece of cable split into an arbitrary number of *segments* (compartments) indexed by *n*. Morphologies with branch points must therefore be represented by more than one section. We here denote the relative length from start to end point of each section by *χ ϵ* [0, 1], see Fig 3A. All segments within the morphology except the initial segment of the *root* section (typically the somatic section) have a *parent* segment indexed by *f*. Each segment in a section can have an arbitrary number of *child* segments, thus a parent segment is the segment which connects to the start point of a *child* segment. We also distinguish between start-, mid-and end-point coordinates of each segment (Fig 3A).

**Figure 3:**
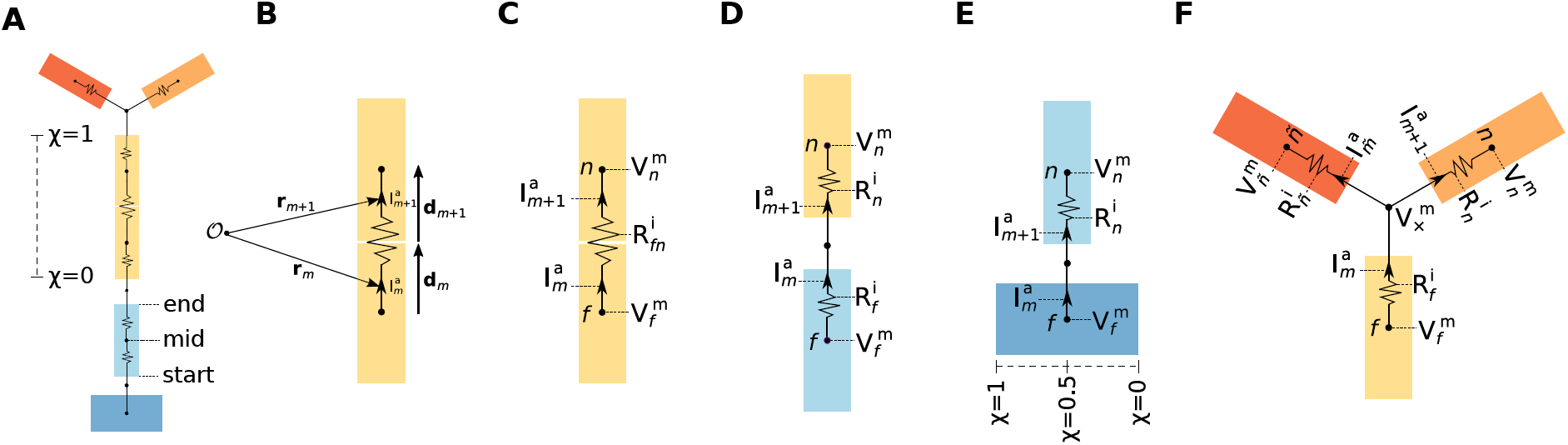
Axial currents in multicompartment neuron models. **A** Schematic illustration of sections (colored rectangles), segments and equivalent electric circuit of a simplified multicompartment neuron model. The relative length *χ* varies between 0 and 1 from start-to end-point of each section. **B** Axial current line element vectors (**d***_m_*, **d**_*m*+1_) and corresponding midpoints (**r***_m_*, **r**_*m*+1_) of axial currents 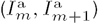 between two connected segments. **C** Axial currents 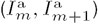, membrane potentials 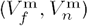, and axial resistance 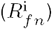 in equivalent electric circuit for a parent segment *f* and child segment *n* in a single section. **D** Similar to panel B, but parent and child segments belong to two different sections. The total series resistance is here *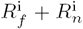*. **E** Illustration of the case where the child segment *n* is connected to a point *χ* = 0.5 on the parent section. For children connected at *χ ϵ* ⟨ 0, 1⟩ the voltage difference 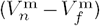 is only across the child segment axial resistance *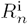*, but the (virtual) current from the node connecting the child start point to the parent midpoint *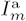* is still accounted for. **F** Illustration of axial currents at branch point between different sections of the morphology. The child segment *n* has one parent *f* and one sibling indexed by 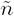, where *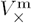* denotes the virtual membrane potential at the node connecting the parent end-point to the children start-points. *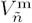* is the voltage in the midpoint of the sibling segment, while 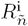 and *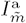* denotes the axial resistance and current between the sibling midpoint and the branch point.

In Fig 3B and C we illustrate the simplest possible calculation of axial current between the midpoints of two neighboring segments *f* and *n* belonging to the same section. Their corresponding membrane voltages are *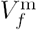* and *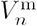*, separated by a total (series) axial resistance *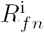*. From NEURON we can easily obtain the axial resistance between the segment midpoint and the segment’s parent node. The parent node is here the midpoint of the parent segment, as the child and parent belong to the same section. Therefore, NEURON gives us the total axial resistance *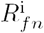* directly, in this case. The axial current magnitude between segment midpoints is then trivial to compute using Ohm’s law (Eq (3)), but as the currents flowing within segments *f* and *n* may not lie on the same axis, we differentiate between the current magnitudes *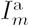* and *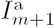*, their axial line element vectors d_*m*_ and d_*m*+1_, and the midpoints of each **r**_*m*_ and **r**_*m*+1_ (panel B). The corresponding current indices are denoted by *m* and *m* + 1 as detailed in Algorithm 1.

Panel D represents the case where the parent and child segments *f* and *n* belong to different sections. The child segment is here the *bottom segment* in a section, and it is connected to the end point of *f*. As the parent node (the node the child segment connects to on the parent segment) is here located between the two segments, NEURON does in this case not give us the total axial resistance directly. Instead, the total (series) axial resistance *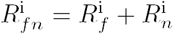* must first be computed to estimate the axial current. *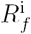* is here the resistance between the parent midpoint and the connecting node, and *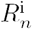* the resistance between the parent node and the segment midpoint.

NEURON allows child sections to be connected anywhere along the parent section: 0 ≤ *χ* ≤ 1. Illustrated in panel E, a child segment is connected to the point *χ* = 0.5 and the axial resistance in the parent segment does not enter the calculation of axial current magnitude. LFPy2.0 still accounts for a virtual axial current *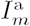* from the parent mid point to the child start point. These virtual currents ensure that the total current dipole moments computed either from transmembrane currents or from axial currents are identical (see Section 2.3.1 for details).

At morphology branch points, several child segments may protrude from a parent segment as illustrated in panel F. As the segment *n* and its sibling 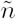 both share the same parent *f*, we estimate the potential *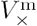* at the branch node using Ohm’s law and Kirchhoff’s current law, accounting for the axial resistivities 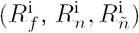 and potentials 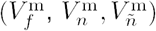, in order to compute the corresponding axial currents *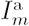* and *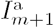*. The full procedure presently used for computing axial currents in LFPy2.0 for the cases illustrated in panels B–F is provided in full detail in Algorithm 1.

##### Algorithm 1 Axial current calculations in LFPy2.0s

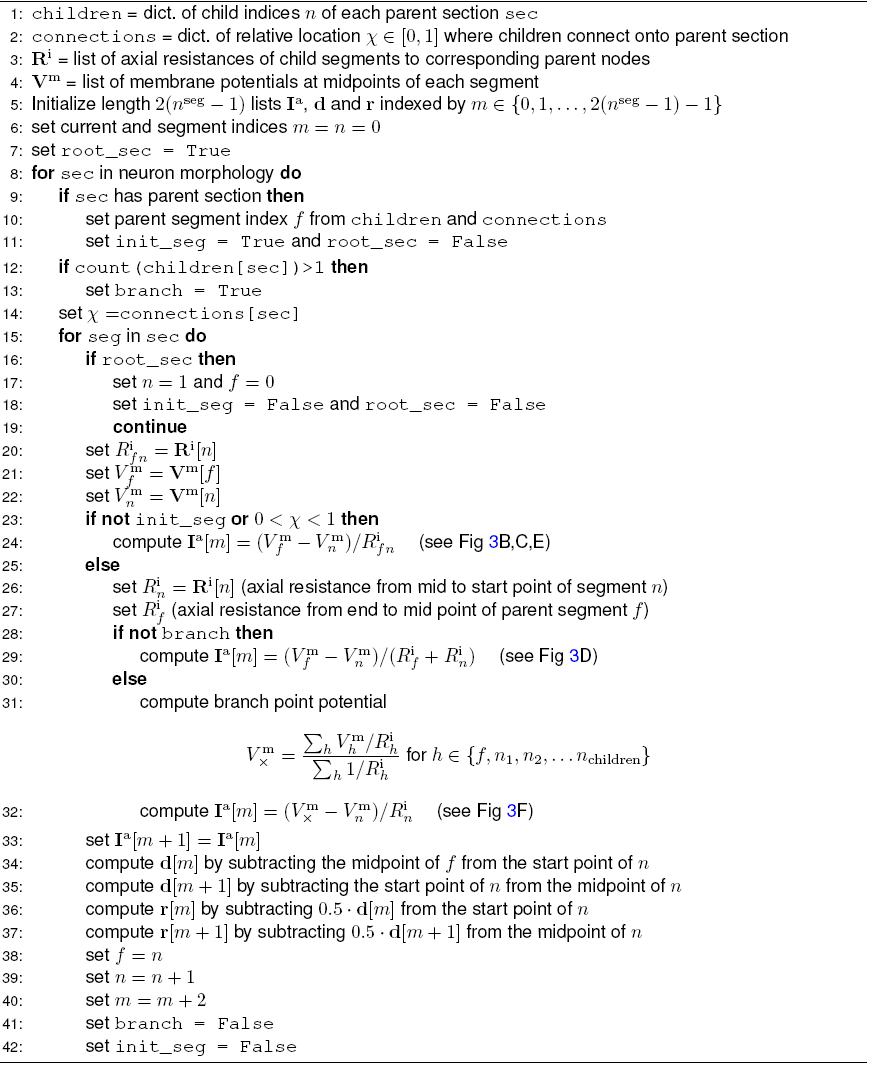

### 2.2 Forward modeling of LFP and MUA signals

The relation between transmembrane currents and extracellular potentials is calculated based on volume conduction theory [Nunez and Srinivasan, 2006; Einevoll et al., 2013]. At the relatively low frequencies relevant in neurophysiology (below a few thousand hertz), this derivation is simplified by omitting terms with time derivatives in Maxwell’s equations (quasistatic approximation, Hämäläinen et al. [1993, p. 426]). Further, the extracellular medium is in all situations considered below assumed to be ohmic, that is, linear and frequency-independent [Pettersen et al., 2012; Einevoll et al., 2013; Miceli et al., 2017].

#### 2.2.1 Homogeneous and isotropic media

We first consider the simplest situation, where the medium is *homogeneous*, i.e., the same in all positions corresponding to an infinite volume conductor, and *isotropic*, i.e., the same electrical conductivity in all directions. The medium is then represented by a scalar extracellular conductivity *σ*_e_. The extracellular potential *φ*(**r**, *t*) at position **r** and time *t* is then given by [Nunez and Srinivasan, 2006; Lindén et al., 2014]

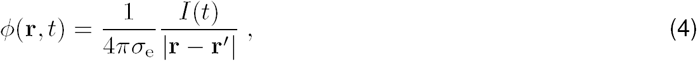

where *I*(*t*) represents a time-varying point current source at position r′. For transmembrane currents 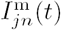 of individual compartments 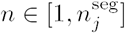 of all cells *j* in a population of *N* cells, the extracellular potential can be computed as the linear sum of their contributions as

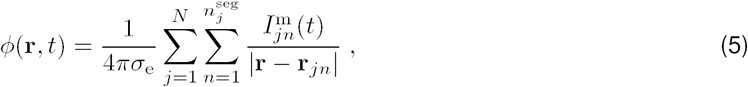

but only under the assumption that each transmembrane current can be represented as a discrete point in space. This point-source assumption can be used in LFPy by supplying the keyword argument and value `method=“pointsource”` to the `RecExtElectrode` class [Lindén et al., 2014].

As a homogeneous current distribution along each cylindrical compartment is assumed, we may employ the *line-source* approximation for somatic and dendritic compartments [Holt and Koch, 1999]. The formula is obtained by integrating Eq (4) along the center axis of each cylindrical compartment *n*, and by summing over contributions from every *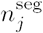* compartment of all *N* cells [Holt and Koch, 1999; Pettersen and Einevoll, 2008; Lindén et al., 2014]:

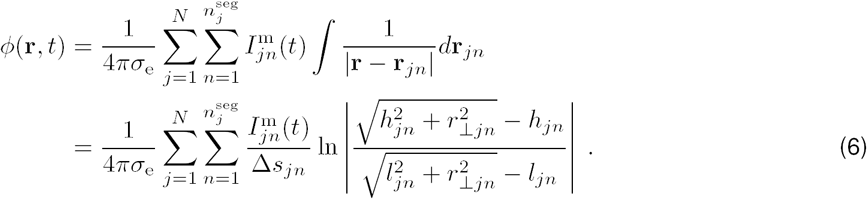

Compartment length is denoted Δ*s_jn_*, perpendicular distance from the electrode point contact to the axis of the line compartment is denoted *r⊥_jn_*, longitudinal distance measured from the start of the compartment is denoted *h_jn_*, and longitudinal distance from the other end of the compartment is denoted *l_jn_* = Δ*s_jn_* + *h_jn_*. The corresponding keyword argument and value to class `RecExtElectrode` is `method=“linesource”` [Lindén et al., 2014].

A final option in LFPy is however to approximate the typically more rounded soma compartments as spherical current sources, thus the line-source formula (Eq (6)) for dendrite compartments is combined with the point-source equation (Eq (4)), obtaining [Lindén et al., 2014]:

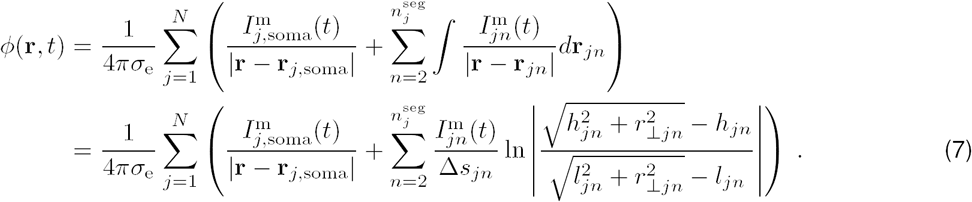

The corresponding keyword argument and value is `method=“soma_as_point”`.

If the distance between current sources and electrode contacts is smaller than the radius of the segment, unphysical singularities may occur in the computed extracellular potential. Singularities are in LFPy automatically prevented by either setting *r⊥*_*jn*_ or |**r** − **r**_*jn*_| equal to the cylindrical compartment radius dependent on the choice of line or point sources.

Electrode contacts of real recording devices have finite spatial extents. A good approximation to the electric potential across the uninsulated surface of metal electrode contact is obtained by computing the spatially averaged electric potential [Robinson, 1968; Nelson et al., 2008; Nelson and Pouget, 2010; Ness et al., 2015], in particular for current sources being located at distances larger than approximately one electrode radius [Ness et al., 2015]. The *disc-electrode* approximation to the potential [Camuñas-Mesa and Quiroga, 2013; Lindén et al., 2014; Ness et al., 2015]

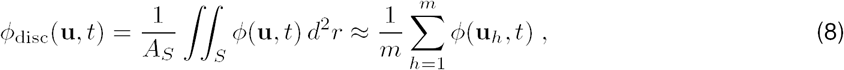

is incorporated in LFPy, with corresponding parameters for contact radius *r*_contact_, number *m* of random points **u**_*h*_ on the flat, circular electrode contact surface when averaging [Lindén et al., 2014]. The surface normal vector for each electrode contact must also be specified.

#### 2.2.2 Discontinuous and isotropic media

Above we described the case for an infinite volume conductor, that is, a constant extracellular conductivity *σ*_e_, as implemented in the initial LFPy release [Lindén et al., 2014]. For cases where *σ*_e_ vary with position, i.e., *σ*_e_ = *σ*_e_(**r**), such as for cortical *in vivo* recordings close to the cortical surface [Einevoll et al., 2007] or *in vitro* recordings using microelectrode arrays (MEAs) [Ness et al., 2015], this approximation does not generally hold. Instead a generalized Poisson equation must be solved [Nicholson and Freeman, 1975]:

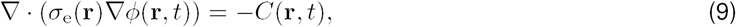

where *C*(**r**, *t*) is the current-source density. This equation can always be solved numerically by means of the Finite Element Method (FEM) [McIntyre and Grill, 2001; Ness et al., 2015] or other mesh-based methods (see for example Tveito et al. [2017]).

In the special case where the conductivity *σ*_e_ is discontinuous in a single direction, that is, a constant con-ductivity in the *xy*-plane and a piecewise constant *σ*_e_(*z*) in the *z*-direction, the ‘Method-of-Images’ (MoI) can be used to make analytical formulas for the extracellular potentials, analogous to Eq (4)–Eq (7) above [Nicholson and Llinas, 1971; Nunez and Srinivasan, 2006]. When applicable, these formulas substantially simplify the modeling of the extracellular potentials compared to FEM modeling.

##### Electrical potentials across microelectrode arrays (MEAs)

The first MoI application is to model recordings in a MEA setting where a slice of brain tissue is put on an insulating recording chip (MEA-chip) and covered with saline [Ness et al., 2015; Hagen et al., 2015]. In this three-layer situation separate conductivity values are assigned to the topmost saline layer conductivity *σ*_S_ for *z ϵ* [*h, ∞*], the middle tissue layer conductivity *σ*_T_ for *z ϵ* [0, *h*) and the lowermost electrode *σ*_G_ for *z ϵ* [–*∞*, 0). The parameter *h* denotes the thickness of the middle tissue layer. The corresponding implementation is provided by the class `RecMEAElectrode`, and has at present the limitations that all current sources (segments) must be contained on the interval *z ϵ* [0, *h*), and that the line-source approximation can only be used when *σ*_G_ = 0 and when computing extracellular potentials for *z* = 0. For other forward-model configurations (for example for 0 ≤ *z* ≤ *h* and/or *σ*_G_ > 0) the point-source approximation can be used. For a detailed derivation of the MoI with two planar electrical boundaries, see Eq. (4) in Ness et al. [2015]. A corresponding example is provided with LFPy2.0 (`example_MEA.py`) which illustrates the computation of extracellular potentials as recorded by a MEA following synaptic activation of a pyramidal cell model.

##### Electrical potentials close to cortical surface

The second MoI application is to model *in vivo* recordings of electrical potentials at or immediately below the cortical surface, that is, the interface between cortical gray matter and dura. Here the extracellular conductivity above the cortical surface *σ*_S_ can be higher or lower than the conductivity in cortical gray matter *σ*_T_ depending on how the measurements are done, for example whether saline or oil is used to cover an inserted laminar electrode [Einevoll et al., 2007]. Such a conductivity jump will affect both the electrical potential recorded at the cortical surface (ECoG recording) as well as the potentials recorded in the top cortical layers [Pettersen et al., 2006]. This can be modeled with the same framework as above, that is, by using the class `RecMEAElectrode`, with the cortical surface at height *h*, while ignoring the lower planar boundary by setting *σ*_G_ = *σ*_T_. In this situation the potential at or below the cortical surface at position (*x, y, z*) for a current source, *I*(*t*), positioned at (*x*′, *y*′, *z*′) is given by [Pettersen et al., 2006; Nunez and Srinivasan, 2006; Ness et al., 2015] as:

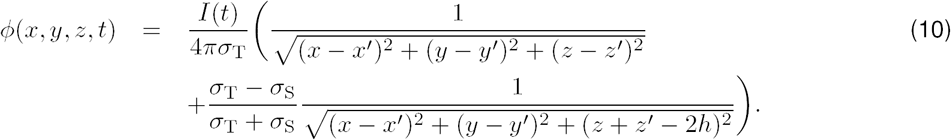

This approach assumes a flat cortical surface. Note, however, that in LFPy2.0 the ECoG signal can also be modeled by means of the four-sphere EEG head model as described below in Sec. 2.3.4. An example is provided with LFPy2.0 (`example_ECoG.py`) which illustrates extracellular potentials recorded in the cortex and at the cortical surface following activation of multiple synapses distributed across a pyramidal cell model.

##### Electrical potentials in spherical conductor

LFPy2.0 also incorporates a spherical conductor model, adapted from [Deng, 2008], where the conductivity is constant within the sphere and zero outside (class `OneSphereVolumeConductor`). Note that this model is applicable for monopolar current sources, unlike the more complex multi-sphere head models described below in Section 2.3 which only apply to dipolar current sources. Although not pursued here, one application of this volume-conductor model could possibly be modeling of LFPs measured in spheroidal brain nuclei.

#### 2.2.3 Homogeneous and anisotropic media

For homogeneous media, i.e., when the extracellular conductivity is the same at all positions, we also added support for anisotropic media [Nicholson and Freeman, 1975]. In this case the extracellular conductivity in Eq (9) must be replaced by a rank 2 (3×3) tensor where the diagonal elements are *σ*_*x*_, *σ*_*y*_, and *σ*_*z*_ and the off-diagonal elements are zero [Nicholson and Freeman, 1975]. This could for example be used to mimic experimental observations of such anisotropy in cortex [Goto et al., 2010], that is, electric currents flow with less resistance along the depth direction (*z*-direction) than in the lateral directions (*x, y*-directions). In this case *σ*_*z*_ > *σ*_*x*_ = *σ*_*y*_ [Ness et al., 2015]. The corresponding implementation is based on the description and implementation provided by Ness et al. [2015], and is in LFPy presently supported by the class `RecExtElectrode`, but not the class `RecMEAElectrode`.

### 2.3 Forward modeling of EEG, ECoG, and MEG signals from current dipoles

The forward modeling of EEG and MEG signals from current dipoles has a long history [Hämäläinen et al., 1993; Nunez and Srinivasan, 2006]. Here the EEG contacts and the MEG magnetometers are located so far away from the neural sources that only the current dipole moments contribute to the measured signals, that is, the contributions from higher-order current multipoles are negligible. From charge conservation, it follows that current monopoles do not exist. To compute the contribution to EEG and MEG signals from detailed neuron models, we thus first need to compute single-neuron current dipole moments, cf. Sec. 2.3.1. Next these must be combined with appropriate volume-conductor models for the head.

In LFPy2.0 we include two ‘head’ models for computing EEG signals from current dipole moments: the (very simplified) infinite homogenous volume-conductor model (Sec. 2.3.2), and the much more involved four-sphere head model where the brain tissue, cerebrospinal fluid (CSF), skull and scalp are represented with different values for the electrical conductivity [Nunez and Srinivasan, 2006; Næss et al., 2017], cf. Sec. 2.3.3. For the MEG signals the forward model is simpler as the magnetic permeability is the same throughout the head as in free space [Hämäläinen et al., 1993]. In LFPy2.0 we include simulation code for computing neural contributions to MEG signals applicable for all head models with spherically-symmetric electrical conductivities, for example, the four-sphere head model, cf. Sec. 2.3.5. While these head models allow for direct calculation of EEG and MEG signals from neurons, it should be noted the computed current dipole moments also can be used for subsequent calculation of EEG and MEG signals by means of boundary element (BEM) or finite element models (FEM) with anatomically detailed head models [DeMunck et al., 2012; Bangera et al., 2010; He et al., 2002; Huang et al., 2016].

#### 2.3.1 Calculation of current dipole moments

##### Current dipole moments from transmembrane currents

The current dipole moment from a single neuron can be computed from transmembrane currents as [Lindén et al., 2010]:

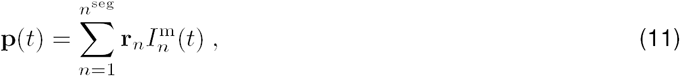

where *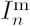* is the transmembrane current at time *t* from compartment *n* at position **r***_n_*. For a population of *N* cells with *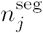* compartments each, the current dipole moment at discrete time steps can be formulated as the matrix product:

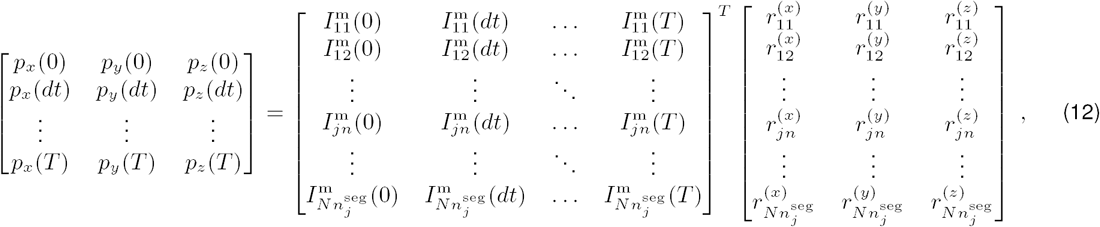

where *pu*(*t*) is the *u*-component (*u ϵ* {*x, y, z*}) of the current dipole moment at time *t* (thus 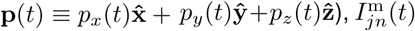 the transmembrane currents of segment *n* of cell *j* at time *t* and *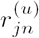* the corresponding *u*-coordinates of each segment’s midpoint. 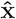, 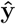 and 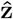 denote the cartesian unit vectors. For more compact notation we here show the transpose (denoted by the raised *T*) of the matrix containing transmembrane currents. Note that the same formula may be used to also compute current dipole moments **p***j* of individual cells *j* (or subsets thereof) by slicing the corresponding matrix elements.

##### Current dipole moments from axial currents

Alternatively, the current dipole moment can be computed from axial currents between neighboring segments (see Section 2.1.2). As an example, we consider a two-compartmental dendritic stick model, where segment *one* will act as a current sink, and segment *two* as a current source. The transmembrane current entering segment two *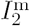* will be the same as the axial current *I*^a^ between the two segments, which is also equal to the current leaving compartment one *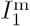*, such that 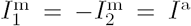. An axial line element vector **d** represents the path traveled by the axial current, which corresponds to the displacement **r**_1_-**r**_2_ between the compartment midpoints. From Eq (11) it thus follows that the current dipole moment is:

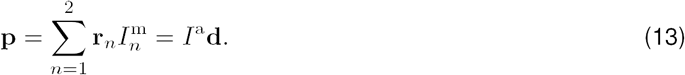

Multiplying each axial current with the respective current path gives a set of current dipoles:

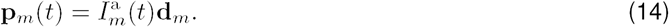

Calculating sets of current dipole moments from neural simulations can be useful, for example for ECoG predictions (see Section 2.3.4) or magnetic fields in proximity of the neuron (see Section 2.4).

#### 2.3.2 EEG signal for homogeneous volume conductor

From eletrostatic theory we have that the electric potential outside a spatial distribution of current sinks and sources can be described by a multipole expansion *φ*(*r*) = *C*_monopole_*/R* + *C*_dipole_*/R*^2^ + *C*_quadrupole_*/R*^3^ + *C*_octupole_*/R*^4^ + *…*, where *R* is the relative distance from the multipole to measurement location (and the coefficients *C* depends on the spherical angles). Due to charge conservation, current opoles do not exist [Nunez and Srinivasan, 2006]. For sufficiently large values of *R* where *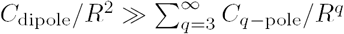*, the electric potential of a neuron can be approximated solely from its current dipole moment, as contributions from quadrupolar and higher-order terms become negligible. The electric potential from a current dipole in an ohmic, homogeneous and isotropic medium is given by [Nunez and Srinivasan, 2006]

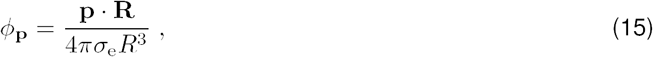

where **p** is the current dipole moment as defined above, *σ*_e_ the conductivity of the extracellular medium, **R** = **r − r′** the displacement vector between dipole location **r′** and measurement location **r**, and *R* = |**R|**. Predictions of extracellular potentials from current dipole moments in homogeneous media are provided by the class `InfiniteVolumeConductor`.

#### 2.3.3 EEG signal in four-sphere head model

The computation of EEG signals assuming a homogeneous volume conductor model is obviously a gross approximation as it neglects the large variation in the extracellular conductivity in the head. In order to com-pute more realistic EEG signals from underlying neuronal sources, we implemented in LFPy2.0 the inhomogeneous four-sphere head model in class `FourSphereVolumeConductor`. This model is composed of four concentric shells representing brain tissue, cerebrospinal fluid (CSF), skull and scalp, where the conductivity can be set individually for each shell [Srinivasan et al., 1998; Nunez and Srinivasan, 2006]. Note that corrections to the original model formulation was recently provided in Næss et al. [2017].

The analytical model solution takes different forms for radial and tangential dipoles. The radial dipole contribution to extracellular potential can be calculated as follows [Næss et al., 2017]:

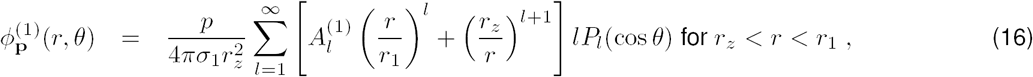

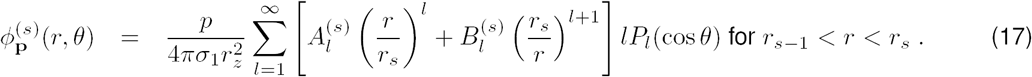

The tangential dipole contribution to the extracellular potentials is:

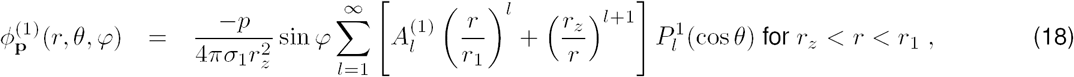

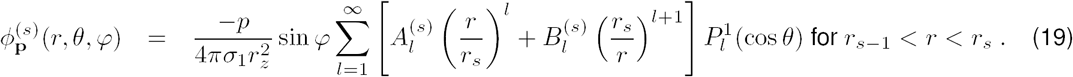

Here, *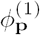* is the extracellular potential measured at radial location *r* in the inner sphere, the brain, while *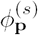* gives the potential in CSF, skull and scalp with *s ϵ* {2, 3, 4}, respectively.

The current dipole moment has magnitude *p* = |**p**| and radial location *r*_*z*_, while *r*_*s*_ and *σ*_*s*_ denote the external radius and conductivity of shell *s*. *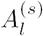* and *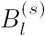* are coefficients that depend on the shell radii and conductivities. *P*_*l*_ (cos *θ*) is the *l*-th Legendre Polynomial where *θ* is the angle between the measurement and dipole location vectors, further *φ* is the azimuth angle and *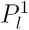* is the associated Legendre polynomial. The detailed derivations of the constants *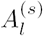* and *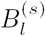* are given in Næss et al. [2017]. Here, we use the notation *σ*_*ij*_ = *σ*_*i*_/*σ*_*j*_ and *r*_*ij*_ ≡ *r*_*i*_/*r*_*j*_:

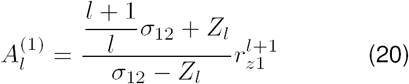

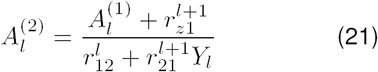

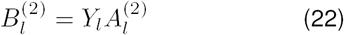

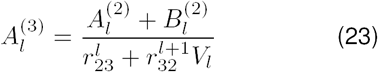

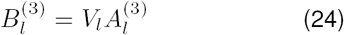

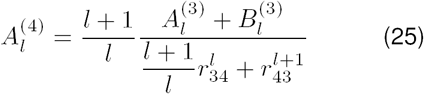

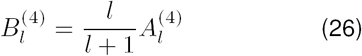

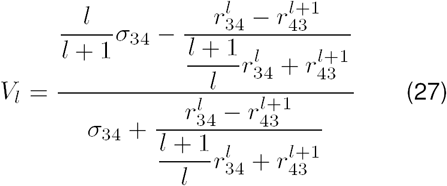

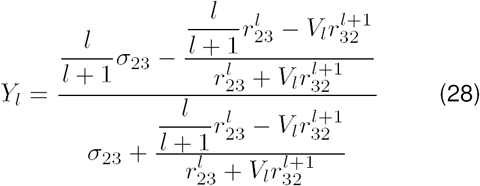

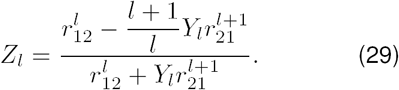

#### 2.3.4 ECoG signal from four-sphere head model

The four-sphere head model is not restricted to EEG predictions, but can also be applied for modeling electric potentials in other layers of the inhomogeneous head model, such as ECoG signals at the interface between the brain tissue and the CSF. In contrast to EEG electrodes, however, the ECoG electrodes are located only micrometers away from the apical dendrites. The electrode’s proximity to the neuronal source makes the four-sphere model a less obvious candidate model, as the model is based on the current dipole approximation, giving good predictions only when the measurement point is more than some dipole lengths away from the source [Lindén et al., 2010]. However, in the `FourSphereVolumeConductor` class method `calc_potential_from_multi_dipoles()`, this problem can be avoided by taking advantage of the fact that electric potentials sum linearly in ohmic media: Instead of computing a single current dipole moment for the whole neuron, we compute multiple current dipole moments, one for each axial current, as described in Section 2.3.1. Since these current dipoles have small enough source separations for the current dipole approximation to be applicable, we can compute the ECoG signal contribution from each current dipole moment separately, using the four-sphere model. The ECoG signal is finally predicted by summing up each contribution. The corresponding LFPy2.0 example file is `/examples/example_ECoG_4sphere.py`.

#### 2.3.5 MEG signals in spherically-symmetric head models

For spherically-symmetric head models the MEG signal can be computed from the current dipole moments set up by intracellular axial currents [Hämäläinen et al., 1993, p. 428]. To compute magnetic fields **B**_**p**_ from current dipole moments we incorporated the special form of the magnetostatic Biot-Savart law (where magnetic induction effects are neglected) [Nunez and Srinivasan, 2006, Appendix 16] given as:

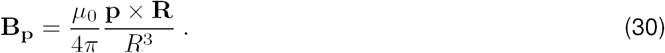

As above, **p** is the dipole source, **R** = **r–r′** the displacement between dipole location **r′** and measurement location **r**, and *R* = |**R|**. For a detailed derivation of this expression see Hämäläinen et al. [1993]. The magnetic field **B** is related to the commonly used quantity **H** (often also termed magnetic field) through **B** = *µ*_0_**H** + **M** = *µ***H** where **M** is the magnetization and *µ* the magnetic permeability of the material. However, in biological tissues the magnetization **M** is very small, and *µ* is very close to the magnetic constant (i.e., the magnetic permeability of vacuum) *µ*_0_ [Hämäläinen et al., 1993]. Predictions of magnetic signals are in LFPy2.0 incorporated in the class `MEG`, which provides the method `calculate_H` in order to compute the magnetic field from a current dipole moment time series. Its output must be multiplied by *µ* to obtain the magnetic field **B**_**p**_.

Throughout this paper, we show for the four-sphere head model magnetic field components decomposed into tangential and radial components at different positions on spherical surfaces. The tangential components were computed in the direction of the angular unit vectors 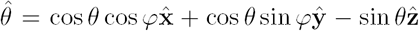 and 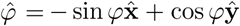 as **B** · 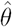 and **B** · 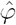, respectively. The radial component was computed as **B**_p_ · 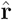 where 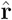 denotes the radial unit vector from the center of the sphere in the direction of the contact. Furthermore, we also show tangential and radial components of the surface magnetic field where the underlying dipoles were rotated by an angle *θ* = *π/*2 around the *x*-axis, denoted 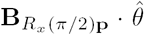 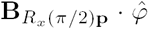 and 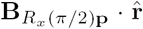, respectively. For this purpose we used the rotation matrix

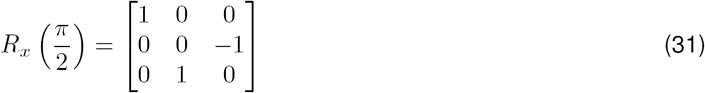

multiplied with the current dipole moment **p** in cartesian coordinates.

### 2.4 Magnetic signals close to neurons

Most studies of magnetic fields generated by neural activity have been based on MEG recordings where the neuronal sources are so distant from the magnetic-field sensors that the far-field dipole approximation in Eq (30) can be applied. However, probes are also being developed for measuring magnetic fields in direct vicinity of the neurons [Barbieri et al., 2016; Caruso et al., 2017]. To compute the magnetic fields in the vicinity of neurons, LFPy2.0 also implements the relevant Biot-Savart law for this situation [Blagoev et al., 2007]:

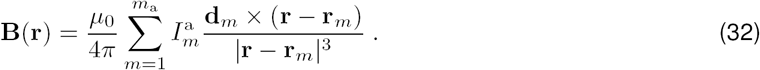

This formula provides the magnetic field for *m*_a_ axial currents *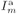* where **d***m* are axial line element vectors, and **r**_*m*_ the midpoint positions of each axial current. The use of this formula assumes that contributions to the magnetic fields from extracellular volume currents are negligible [Hämäläinen et al., 1993, p. 427]. Predictions of magnetic signals from axial currents (or equivalently sets of current dipoles) are in LFPy2.0 facilitated by the corresponding class method `MEG.calculate_H_from_iaxial()`. We show (in Fig 2) the *y*-components of the magnetic fields in vicinity of a model neuron computed as **B** · ŷ and **B**_p_ · ŷ respectively.

### 2.5 New Classes and network use-case implementation

The first release of LFPy described in Lindén et al. [2014] included a set of Python class definitions for instantiating single-cell models (`Cell, TemplateCell`) and corresponding instrumentation of the models with synapse point processes attached to the cell (`Synapse`), patch-clamp electrodes (StimIntElectrode) and extracellular recording electrodes (`RecExtElectrode`). Simulations with multiple simultaneous cell-object instances were at the time not supported. Class `TemplateCell` supported the use of template specifications, a requirement for networks in NEURON, but was primarily written to support source codes of ‘network-ready’ single-cell models such as the Hay et al. [2011] models of layer-5 pyramidal neurons available from, for example, ModelDB (senselab.med.yale.edu/modeldb, McDougal et al. [2017]).

The ‘one cell at a time’ approach may seem limited, in particular when considering ongoing network interactions, but knowing that forward-modeling of extracellular potentials can be decoupled from the network simulation, users could always set up simulations of each individual cell, play back synapse activation times as occurring in the connected network, and sum up the single-cell contributions to the extracellular potential. Thus, the calculation of extracellular potentials can even be dealt with in an ‘embarrassingly’ parallel manner [Foster, 1995; Hagen et al., 2016]. These simplifying steps are not possible if the extracellular potential itself affects the cellular dynamics, that is, if mutual interactions between cellular compartments belonging to the same or different cells occur through the extracellular potential, so-called ephaptic interactions [Anastassiou et al., 2011; Goldwyn and Rinzel, 2016; Tveito et al., 2017].

For the present LFPy2.0 release, we added support for simulations of recurrently connected multicompartment models with concurrent calculations of extracellular potentials and current dipole moments. As described above, the current dipole moment is used for predictions of distal electric potentials (for example scalp surface potentials as in EEG measurements) and magnetic fields (as in MEG measurements). For our example use case, we considered a recurrent network of four populations of multicompartment neuron models. We added a new set of generic class definitions in LFPy to represent the network, its populations and neurons, as well as classes representing different volume-conductor models and measurement modalities as summarized next.

#### Cells

Each individual neuron in an LFPy network exists as an instantiation of class `NetworkCell`. As this class definition uses class inheritance from the old `TemplateCell` and in turn `Cell` classes, it retains all common methods and attributes from its parent classes. The `NetworkCell` can therefore be instantiated in a similar manner as its parent class:

~~~
#!/usr/bin/env python “““example_NetworkCell.py”““ # import modules:
**from** LFPy **import** NetworkCell, StimIntElectrode
**from** matplotlib.pyplot **import** subplot, plot
# class NetworkCell parameters:
cellParameters = dict(
    morphology=‘BallAndStick.hoc’,
    templatefile=‘BallAndStickTemplate.hoc’,
    templatename=‘BallAndStickTemplate’,
    templateargs=None,
    v_init=-65.
    )
# create cell:
cell = NetworkCell(
    tstart=0., tstop=100.,
    **cellParameters
    )
# create stimulus device: iclamp = StimIntElectrode(
    cell=cell,
    idx=0,
    pptype=‘IClamp’,
    amp=0.5,
    dur=80.,
    delay=10.,
    record_current=True
    )
# run simulation:
cell.simulate()
# plot cell response:
subplot(2,1,1)
plot(cell.tvec, iclamp.i)
subplot(2,1,2)
plot(cell.tvec, cell.somav)
~~~

The morphology and template files referred to above are defined in NEURON ‘hoc’ language files. A ‘ball and stick’ style morphology file with active soma (Hodgkin & Huxley Na^+^, K^+^ and leak channels) and passive dendrite sections and corresponding template file was written as:

~~~
/* -------------------------------
BallAndStick.hoc
-------------------------------*/
// Create sections:
create soma[1]
create apic[1]

// Add 3D information:
soma[0] {
    pt3dadd(0, 0, −15, 30)
    pt3dadd(0, 0, 15, 30)
}
apic[0] {
    pt3dadd(0, 0, 15, 3)
    pt3dadd(0, 0, 1015, 3)
}

// Connect section end points:
connect apic[0](0), soma[0](1)

// Set biophysical parameters:
forall {
    Ra = 100.
    cm = 1.
    all.append()
}
soma { insert hh }
apic {
    insert pas
    g_pas = 0.0002
    e_pas = −65.
}
/* ----------------------------*/
~~~

and

~~~
/* -------------------------------
BallAndStickTemplate.hoc
-------------------------------*/
begintemplate BallAndStickTemplate
public init, soma, apic
public all
objref all proc init() {
    all = new SectionList()
}
create soma[1], apic[1]
endtemplate BallAndStickTemplate
/* ----------------------------*/
~~~

In contrast to class `TemplateCell`, class `NetworkCell` has built-in methods to detect somatic action potentials and set-ups of synapses being activated by such threshold crossings in other cells.

#### Network populations

One step up in the hierarchy, class `NetworkPopulation` represents a size *NX* population of `NetworkCell` objects of one particular cell type (*X*) in the network. The class can be used directly as:

~~~
#!/usr/bin/env python
“““example_NetworkPopulation.py”””
# import modules
**from** mpi4py.MPI **import** COMM_WORLD as COMM
**from** LFPy **import** NetworkPopulation, NetworkCell
# class NetworkCell parameters:
cellParameters = dict(
    morphology=‘BallAndStick.hoc’,
    templatefile=‘BallAndStickTemplate.hoc’,
    templatename=‘BallAndStickTemplate’,
    templateargs=None,
    delete_sections=False,
)
# class NetworkPopulation parameters:
populationParameters = dict(
    Cell=NetworkCell,
    cell_args = cellParameters,
    pop_args = dict(
        radius=100.,
        loc=0.,
        scale=20.),
    rotation_args = dict(x=0., y=0.),
)
# create population:
population = NetworkPopulation(
    first_gid=0, name=‘E’,
    **populationParameters
    )
# print out some info:
**for** cell **in** population.cells:
    **print**(‘RANK {}; pop {}; gid {}; cell {}’.format(
    COMM.Get_rank(), population.name,
    cell.gid, cell))
~~~

Direct instantiation of class `NetworkPopulation`, however, is of limited use as it does not provide any means of simulation control by itself, and has only one built-in method to draw and set random cell-body positions within a chosen radius (`pop_args[‘radius’]`) and depth from the normal distribution 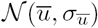. In the code example above, `pop_args[‘loc’]` refers to expected mean depth 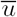 and `pop_args[‘scale’]` to the corresponding standard deviation *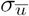*. A random cell rotation around its own vertical *z*-axis is applied by default. The integer `cell.gid` value accessed above is a unique global identifier *g*_id_ of each cell in the network, and is assigned in running order from the number `first_gid`. For parallel execution using MPI, cells will be distributed among threads according to the round-robin rule if the condition *g*_id_%*N*_MPI_ == *k* is True, where % denotes a division modulus operation, *N*_MPI_ the MPI pool size and *k ϵ* [0, 1, *…, N*_MPI_ 1] the corresponding rank number.

##### Networks

The new network functionality is provided through class Network. An instantiation of the class sets attributes for the default destination of file output, temporal duration *t* and resolution *dt* of the simulation, a chosen initial voltage *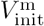* and global temperature control *T*celsius (which affects channel dynamics). Furthermore, the class instance provides built-in methods to create any number of *NX*-sized populations *X*. Different built-in class methods create random connectivity matrices 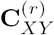 (per rank, see *Connectivity Model* below) between any presynaptic population *X* and postsynaptic population *Y*, and connect *X* and *Y* using an integer number of synapses per connection *n*^syn^ drawn from the capped normal distribution *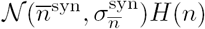* where *H*(*·*) denotes the Heaviside step function. Similarly, synaptic conductances *g*^syn^ are drawn from the distribution *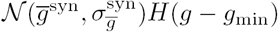* (where *g*_min_ denotes minimum synaptic conductance) with connection delays *δ*^syn^ from *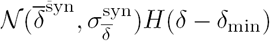* (where *δ*_min_ denotes the minimum delay in the network). The network class handles the synapse model in NEURON and corresponding parameters (time constants, reversal potentials, putative synapse locations etc.), and finally provides a simulation control procedure. The simulation control allows for concurrent calculation of network activity and prediction of extracellular potentials as well as the current dipole moment.

In order to set up a complete network simulation we may choose to define NetworkCell and `NetworkPopulation` parameters as above, and define parameter dictionaries for our instances of Network and extracellular measurement device `RecExtElectrode`:

~~~
#!/usr/bin/env python
“““example_Network.py”””
# import modules
**import** numpy as np
**import** scipy.stats as st
**from** mpi4py**import** MPI
**from** LFPy **import** NetworkCell, Network
**import** neuron
# relative path for simulation output:
OUTPUTPATH=‘example_network_output’
# class NetworkCell parameters:
cellParameters = dict(**cellParameters)
# class NetworkPopulation parameters:
populationParameters = dict(**populationParameters)
# class Network parameters:
networkParameters = dict(
    dt = 2**-4,
    tstop = 1200.,
    v_init = −65.,
    celsius = 6.5,
    OUTPUTPATH = OUTPUTPATH
)
# class RecExtElectrode parameters:
electrodeParameters = dict(
    x = np.zeros(13), y = np.zeros(13),
    z = np.linspace(1000., −200., 13),
    N = np.array([[0., 1., 0.]]*13),
    r = 5.,
    n = 50,
    sigma = 0.3,
)
# method Network.simulate() parameters:
networkSimulationArguments = dict(
    rec_current_dipole_moment = True,
    rec_pop_contributions = True,
)
~~~

Furthermore, we define population names (*X*) and corresponding sizes (*N_X_*), as well as one overall connection probability (*C*_*Y X*_):

~~~
# population names, sizez and connection probability:
population_names = [‘E’, ‘I’]
population_sizes = [80, 20]
connectionProbability = [[0.05, 0.05]]*2
~~~

Then, we may chose to define the synapse model and corresponding parameters (here using NEURON’s built-in two-exponential model Exp2Syn) for synapse conductances (weight), delays and synapses per con-nection (multapses), as well as layer-specificities of connections (*ℒ*_*Y XL*_, see Hagen et al. [2016] and below):

~~~
# synapse model. All corresponding parameters for weights,
# connection delays, multapses and layerwise positions are
# set up as shape (2, 2) nested lists for each possible
# connection on the form:
# [[“E:E”, “E:I”],
# [“I:E”, “I:I”]].
synapseModel = neuron.h.Exp2Syn # synapse parameters
synapseParameters = [[dict(tau1=0.2, tau2=1.8, e=0.)]*2,
                     [dict(tau1=0.1, tau2=9.0, e=-80.)]*2]
# synapse max. conductance (function, mean, st.dev., min.):
weightFunction = np.random.normal
weightArguments = [[dict(loc=0.002, scale=0.0002)]*2,
                   [dict(loc=0.01, scale=0.001)]*2]
minweight = 0.
# conduction delay (function, mean, st.dev., min.):
delayFunction = np.random.normal
delayArguments = [[dict(loc=1.5, scale=0.3)]*2]*2
mindelay = 0.3
multapseFunction = np.random.normal
multapseArguments = [[dict(loc=2., scale=.5)]*2,
                     [dict(loc=5., scale=1.)]*2]
# method NetworkCell.get_rand_idx_area_and_distribution_norm
# parameters for layerwise synapse positions:
synapsePositionArguments = [[dict(section=[‘soma’, ‘apic’],
                               fun=[st.norm]*2,
                               funargs=[dict(loc=500., scale=100.)]*2,
                               funweights=[0.5, 1.])]*2,
                          [dict(section=[‘soma’, ‘apic’],
                               fun=[st.norm]*2,
                               funargs=[dict(loc=0., scale=100.)]*2,
                               funweights=[1., 0.5])]*2]
~~~

Note that we above relied on Python list-comprehension tricks for compactness. Having defined all parameters, one can then create the network, populations, stimulus, connections, recording devices, run the simulation and collect simulation output:

~~~
**if** __name__ == ‘__main__’:
                 ############################################################################
                 # Main simulation
                 ############################################################################
                 # create directory for output:
                 **if not** os.path.isdir(OUTPUTPATH):
                    **if** RANK == 0:
                             os.mkdir(OUTPUTPATH) COMM.Barrier()

# instantiate Network:
network = Network(**networkParameters)

# create E and I populations:
**for** name, size **in** zip(population_names, population_sizes):
       network.create_population(name=name, POP_SIZE=size,
                                **populationParameters)

         # create excitatpry background synaptic activity for each cell
         # with Poisson statistics
         **for** cell **in** network.populations[name].cells:
           idx = cell.get_rand_idx_area_norm(section=‘allsec’, nidx=64)
           **for** i **in** idx:
             syn = Synapse(cell=cell, idx=i, syntype=‘Exp2Syn’,
                           weight=0.002,
                           **dict(tau1=0.2, tau2=1.8, e=0.))
             syn.set_spike_times_w_netstim(interval=200.)

# create connectivity matrices and connect populations:
**for** i, pre **in** enumerate(population_names):
             **for** j, post **in** enumerate(population_names):
                        # boolean connectivity matrix between pre-and post-synaptic neurons
                        # in each population (postsynaptic on this RANK)
                        connectivity = network.get_connectivity_rand(pre=pre, post=post,
                                                       connprob=connectionProbability[i][j])
                       
                        # connect network:
                        (conncount, syncount) = network.connect(
                                    pre=pre, post=post,
                                    connectivity=connectivity,
                                    syntype=synapseModel,
                                    synparams=synapseParameters[i][j],
                                    weightfun=np.random.normal,
                                    weightargs=weightArguments[i][j],
                                    minweight=minweight, delayfun=delayFunction,
                                    delayargs=delayArguments[i][j],
                                    mindelay=mindelay,
                                    multapsefun=multapseFunction,
                                    multapseargs=multapseArguments[i][j],
                                    syn_pos_args=synapsePositionArguments[i][j],
                                    )

# set up extracellular recording device:
electrode = RecExtElectrode(**electrodeParameters)
# run simulation:
SPIKES, OUTPUT, DIPOLEMOMENT = network.simulate(
     electrode=electrode,
     **networkSimulationArguments
)
~~~

The argument `SPIKES` returned by the final `network.simulate` method call is a dictionary with keys gids and `times`, where the corresponding values are lists of global neuron ID’s (*g*ID) and numpy ar-rays with spike times of each respective unit in the network. The returned `OUTPUT` and `DIPOLEMOMENT` arguments are numpy arrays with structured datatypes (sometimes referred to as record arrays). The ar-ray `OUTPUT[‘imem’]` is the total extracellular potential from all transmembrane currents in units of mV, the entries `‘E’` and `‘I’` contributions from the excitatory and inhibitory neuron populations, respectively. DIPOLEMOMENT similarly contains the current dipole moment from populations `‘E’` and `‘I’`, but not the sum as the current dipole moment of different populations may, in principle, be freely positioned in different locations within a volume conductor. The computed current dipole moments by themselves have no well defined positions in space and must explicitly be assigned a position by the user, unlike the individual compartment positions used when computing the extracellular potential.

The corresponding LFPy2.0 example files discussed throughout this section are:

- `/examples/example_network/example_NetworkCell.py,`
- `/examples/example_network/example_NetworkPopulation.py`
- `/examples/example_network/example_Network.py.`

#### 2.6 Description of biophysically-detailed network in example use case

##### Neuron models

Our example network model presented in Results comprised about 5500 biophysically detailed multicompartment neurons obtained from The Neocortical Microcircuit Collaboration (NMC) Portal (https://bbp.epfl.ch/nmc-portal, Ramaswamy et al. [2015]). The NMC portal provides NEURON code for about 1000 different single-cell models as well as connectivity data of a reconstruction and simulation of a rat so-matosensory cortex column [Markram et al., 2015].

For simplicity of this demonstration, we here use only four different single-cell models as shown in Fig 4A for the different network populations. For layers 4 and 5 we chose the most common excitatory cell type and most common inhibitory interneuron cell type, in accordance with statistics of the reconstructed microcircuit of Markram et al. [2015] as provided on the NMC portal. The table in panel A summarizes population names (*X*– presynaptic; *Y* – postsynaptic) which here coincide with morphology type (m), electric type (e), cell model #, compartment count per single-cell model 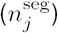, number of cells *N*_*X*_ in each population, occurrence 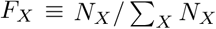 the number of external synapses on each cell *n*_ext_ rate expectation of external synapses *v*_ext_ and the mean *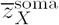 and standard deviation *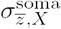* of the normal distribution 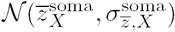* from which somatic depths are drawn for each population. The cell type can be derived from the ‘m’ and ‘e’ type in the table. Using the nomenclature of Markram et al. [2015], L4 and L5 are abbreviations for layer 4 and 5; PC – pyramidal cell; LBC – large basket cell; TTPC1 – thick-tufted pyramidal cell with a late bifurcating apical tuft; MC – Martinotti cell; cAD – continuous adapting; dNAC – delayed non-accommodating; bAC – burst accommodating. Thus, L4_PC_cAD corresponds to a layer 4 pyramidal cell with a continuously adapting firing pattern as a response to depolarizing step current and so forth. As multiple variations of the same cell types are provided on the NMC portal, the cell model # can be used to identify the particular single-cell model and corresponding file sets used in the network described here. These single-cell model files can be downloaded one after another from the portal as for example L5_TTPC1_cADpyr232_1.zip, or all together in a single archive. For simplicity we ignore heterogeneity in e-types for each m-type, thus the population counts *NX* correspond to the count per m-type in the reconstructed microcircuit. Note for the present network description that {*X, Y*, m} ϵ {L4_PC, L4_LBC, L5_TTPC1, L5_MC}.

**Figure 4:**
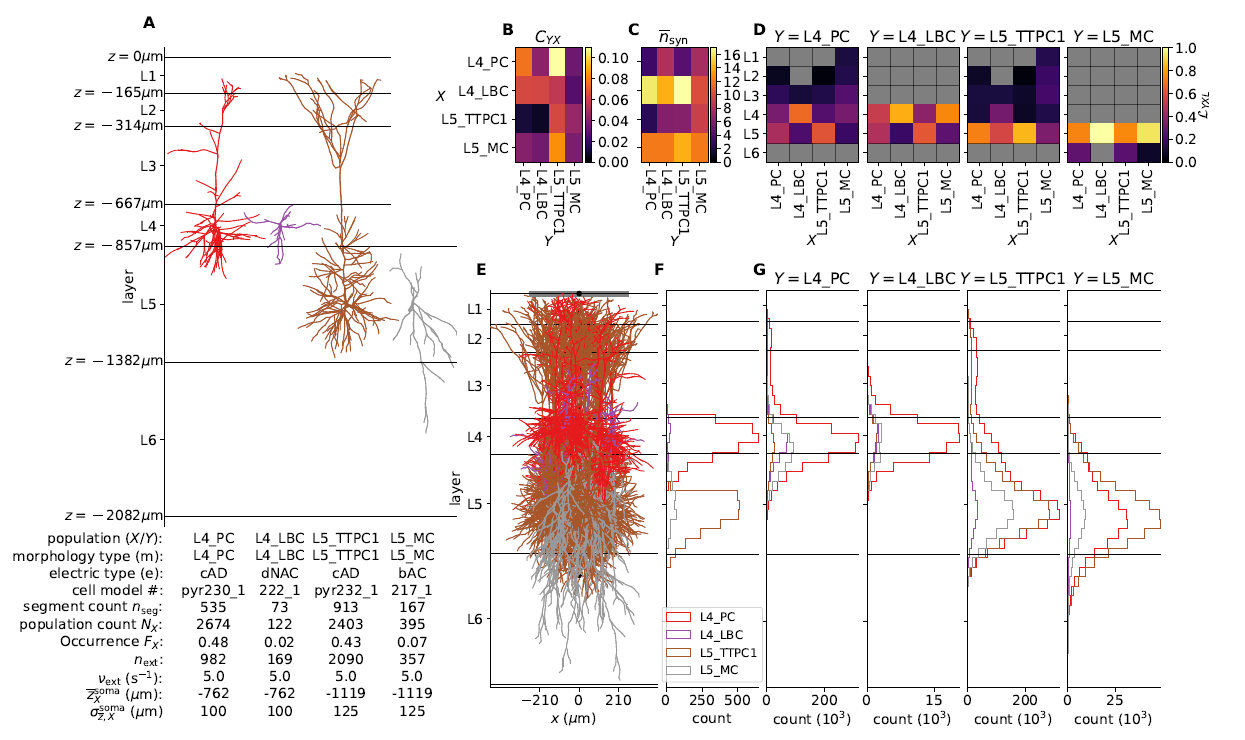
Details of the example network. **A** Biophysically detailed neuron models of the network, with depth-values of boundaries of layers 1-6. The lower left table summarizes population names (*X* – presynaptic; *Y* – postsynaptic) which here coincide with morphology type (m); electric type (e); cell model #; compartment count per single-cell model 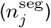; number of cells *N*_*X*_ in each population; occurrence *F_X_* (defined as *N_X_ / Σ_X_ N_X_*); the number of external synapses on each cell *n*_ext_; rate expectation of external synapses *v*_ext_; the expected mean *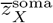* and standard deviation *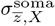* of the normal distribution *𝒩* from which somatic depths are drawn. **B** Pairwise connection probability *C*_*Y X*_ between cells in presynaptic populations *X* and postsynaptic populations *Y*. **C** Average number *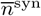* of synapses created per connection between *X* and *Y*. **D** Layer specificity of connections *ℒ*_*Y XL*_ [Hagen et al., 2016] from each presynaptic population *X* onto each postsynaptic population *Y*. Gray values denote *ℒ*_*Y XL*_ = 0. **E** Illustration of cylindrical geometry of populations including a laminar recording device for extracellular potentials (black circular markers) and a single ECoG electrode above layer 1 (gray line). *n* = 20 neurons of each population are shown in their respective locations. **F** Laminar distribution of somas for each network population (Δ*z* = 50 *µ*m) in one instantiation of the circuit. **G** Laminar distribution of synapses across depth onto each postsynaptic population *Y* from presynaptic populations *X* (Δ*z* = 50 *µ*m).

##### Population geometry

The centers of somatic compartments for all cells *i ϵ X* were distributed with even probability within a circular radius of 210 *µ*m corresponding to the radius of the reconstructed somatosen-sory column in Markram et al. [2015]. The corresponding depths were drawn from the normal distribution *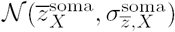* using population-specific mean and standard deviations given in Fig 4A. Neuron positions resulting in any neuron compartments protruding above the hypothetical cortical surface at *z* = 0 or below layer 6 at *z* = *-*2082 *µ*m were redrawn from the depth distribution. All cells were rotated around their local vertical *z*-axis by a random angle *θ ϵ* [0, 2*π*).

##### Synapse models

For synapses made by cells in a presynaptic population *X* onto a postsynaptic population *Y* we used synapse model files provided with the single-cell model files from the NMC portal. There are two base models with connection-specific parameterization which were obtained from the portal. Excitatory synapses are modeled as probabilistic AMPA and NMDA receptors, while inhibitory synapses are modeled as probabilistic GABAA receptors. Both synapse types were modeled with presynaptic short-term plasticity. The synapse parameterization procedure and validation is described in detail in Markram et al. [2015], with code implementations based on Fuhrmann et al. [2002]. The synapse parameters are summarized in Table 1, detailing the synapse model names, average synaptic conductances 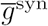 and corresponding standard deviations 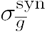, release probabilities *Pu*, relaxation time constants from depression *τ*_Dep_, relaxation time constants from facilitation *τ*_Fac_, ratios of NMDA vs. AMPA (excitatory connections only), rise and decay time constants 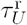 and *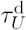* of the two-exponential conductances of each current type *U ϵ* {AMPA, NMDA, GABAA*}*, and reversal potentials *e*^syn^. Random conductances for each individual synapse were drawn from the capped normal distribution *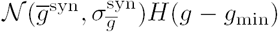*. For our network we set the minimum synaptic conductance to be *g*_min_ = 0 nS.

**Table 1:**
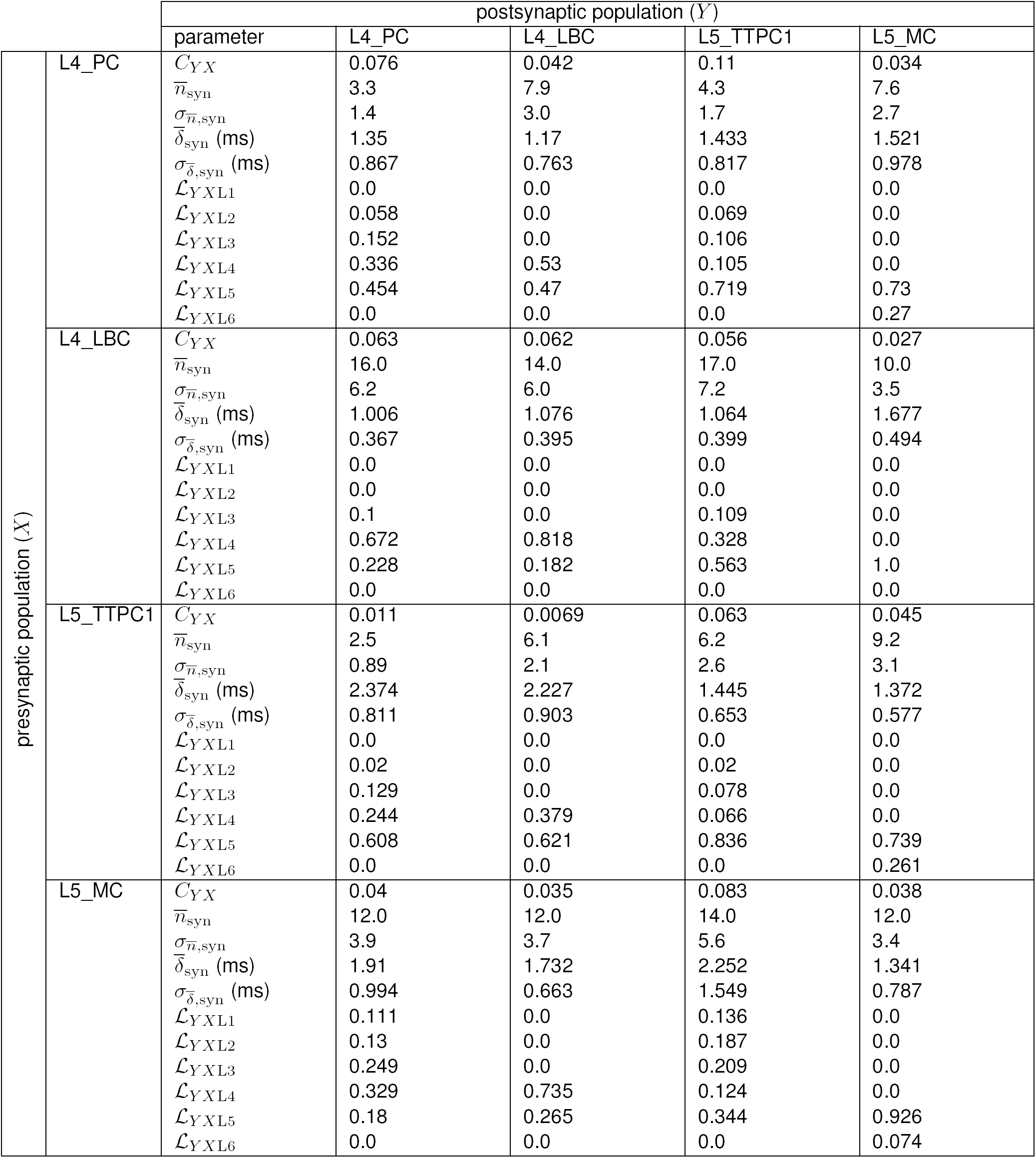
Summary of intrinsic synapse parameters.

##### Extrinsic input

Synapses from external inputs to the neurons in our network were modeled similarly to ex-citatory synapses of intrinsic network connections. For inputs to a population *Y* in layer *L* we chose to duplicate the synapse parameters of connections made by the presynaptic excitatory population within the same layer (as we were unable to assess what parameters were used for extrinsic connections in Markram et al. [2015]). Our synapse parameters are given in Table 2. For each cell in the network we created *n*_ext_ synapses set randomly onto dendritic and apical compartments with compartment specificity of connections*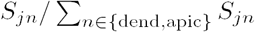*, where *S*_*jn*_ denotes surface area of compartment *n* of cell *j*. The random activation times of each synapse were set using Poisson processes with rate expectation *v*_ext_ for the duration of the simulation. The values for *n*_ext_ and *v*_ext_ are given in Fig 4A, and were set by hand in order to maintain spiking activity in all populations.

**Table 2:** Synapse parameters for extrinsic input.

| | | postsynaptic population ($Y$) | | | | |
| --- | --- | --- | --- | --- | --- | --- |
|  |  | parameter | L4_PC | L4_LBC | L5_TTPC1 | L5_MC |
| presynaptic pop. ($X$) | ext | syn. model | ProbAMPANMDA | ProbAMPANMDA | ProbAMPANMDA | ProbAMPANMDA |
| | | $\bar{g}^{\text{syn}}$ (nS) | 0.3 | 0.33 | 0.31 | 0.33 |
| | | $\sigma_{\bar{g}}^{\text{syn}}$ (nS) | 0.11 | 0.15 | 0.11 | 0.15 |
| | | $P_u$ | 0.859 | 0.254 | 0.5 | 0.252 |
| | | $\tau_{\text{Dep}}$ (ms) | 670 | 700 | 670 | 710 |
| | | $\tau_{\text{Fac}}$ (ms) | 17 | 21 | 17 | 21 |
|  |  | NMDA ratio | 0.4 | 0.4 | 0.4 | 0.4 |
| | | $\tau_{\text{AMPA}}^{\text{r}}$ (ms) | 0.2 | 0.2 | 0.2 | 0.2 |
| | | $\tau_{\text{AMPA}}^{\text{d}}$ (ms) | 8.291 | 8.295 | 8.271 | 8.339 |
| | | $\tau_{\text{NMDA}}^{\text{r}}$ (ms) | 0.29 | 0.29 | 0.29 | 0.29 |
| | | $\tau_{\text{NMDA}}^{\text{d}}$ (ms) | 43 | 43 | 43 | 43 |
| | | $e^{\text{syn}}$ (mV) | 0 | 0 | 0 | 0 |

##### Connectivity model

Random connections in our network were set using the Python-implementation of the ‘connection-set algebra’ of Djurfeldt [2012]; Djurfeldt et al. [2014] (https://github.com/INCF/csa). Using this formalism, we constructed boolean connectivity matrices 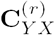 for postsynaptic cells *j*^(*r*)^ *⊂ Y* distributed across each separate parallel MPI rank (denoted by the superset ‘(*r*)’ for rank number) and presynaptic cells *i ϵ X*. Each instance of 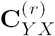 had shape (*NX × Nj*(*r*)*⊂Y*), with entries equal to True denoting connections from cell *i* to *j*^(*r*)^, as expressed mathematically by

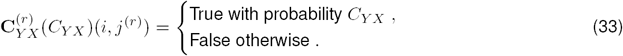

For *X* = *Y* and *i* = *j*^(*r*)^, entries in 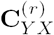 were set to False (no autapses). We used fixed connection probabilities *CY X* as obtained from the NMC portal between our chosen m-types.

##### Multapses

As multiple synapses per connection appear to be a prominent feature in cortical networks (see Reimann et al. [2015]; Markram et al. [2015] and references therein), we drew for every connection between presynaptic cell *i* and postsynaptic cell *j* a random number of synapses *n*^syn^ rounded to an integer from the capped normal distribution *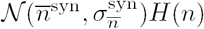*. Conduction delays from action-potential detection (thresh-old *θ*_AP_ = *-*10 mV) in cell *i* for each corresponding synapse onto cell *j* were drawn from the distribution 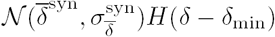. For our network we set the minimum delay *δ*_min_ = 0.3 ms for all connections.

##### Layer-specificity of connections

In order to position each individual synapse of a connection on a cell *j ϵY*, in a simplified manner that depended on the degree of overlap between presynaptic axons and postsynaptic dendrites (‘Peter’s rule’), we calculated for each postsynaptic population *Y* layer-specificities of connections *ℒ*_*Y XL*_ in layer *L* for synapses made by presynaptic populations *X* [Hagen et al., 2016], by first computing the sums Δ*_siXL_* = Σ _*nϵ*axon_ Δ_*sinXL*_, that is, the total axon length of a presynaptic cell type per layer *L* and sums Δ_*sjY L*_ = Σ_*nϵ*{soma,dend}_ Δ_*sjnY L*_ of total dendrite and soma length for each postsynaptic cell type across each layer. Then we defined the layer-specificity of connections as

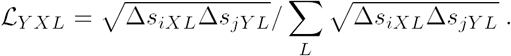

The sums Σ*_L_ ℒ*_*Y XL*_ = 1 for all *X* and *Y*. Synapse sites of connections onto cell *j* were then set randomly with a compartment specificity of connections 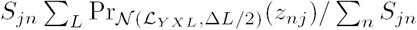, where *S*_*jn*_ is the sur-face area of compartment *n* of the cell *j* centered at depth *z*_*nj*_ and Pr_*N* (*…*)_ the probability density function of the distribution *𝒩* (*ℒ*_*Y XL*_, Δ*L/*2). Δ*L* denotes the thick<u>n</u>ess of layer *L*.

All connectivity parameter values 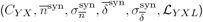 are summarized in Table 3. Visual representations of 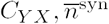 and *ℒ*_*Y XL*_ are shown in Fig 4B, C and D. Panel E shows 20 cells in each population *X* with corresponding distribution of *N*_*X*_ somas across depth (Δ*_z_* = 50 *µ*m) in panel F. Panel G shows the resulting distribution of synapses across depth for all combinations of *Y* and *X* (Δ*_z_* = 50 *µ*m).

**Table 3:**
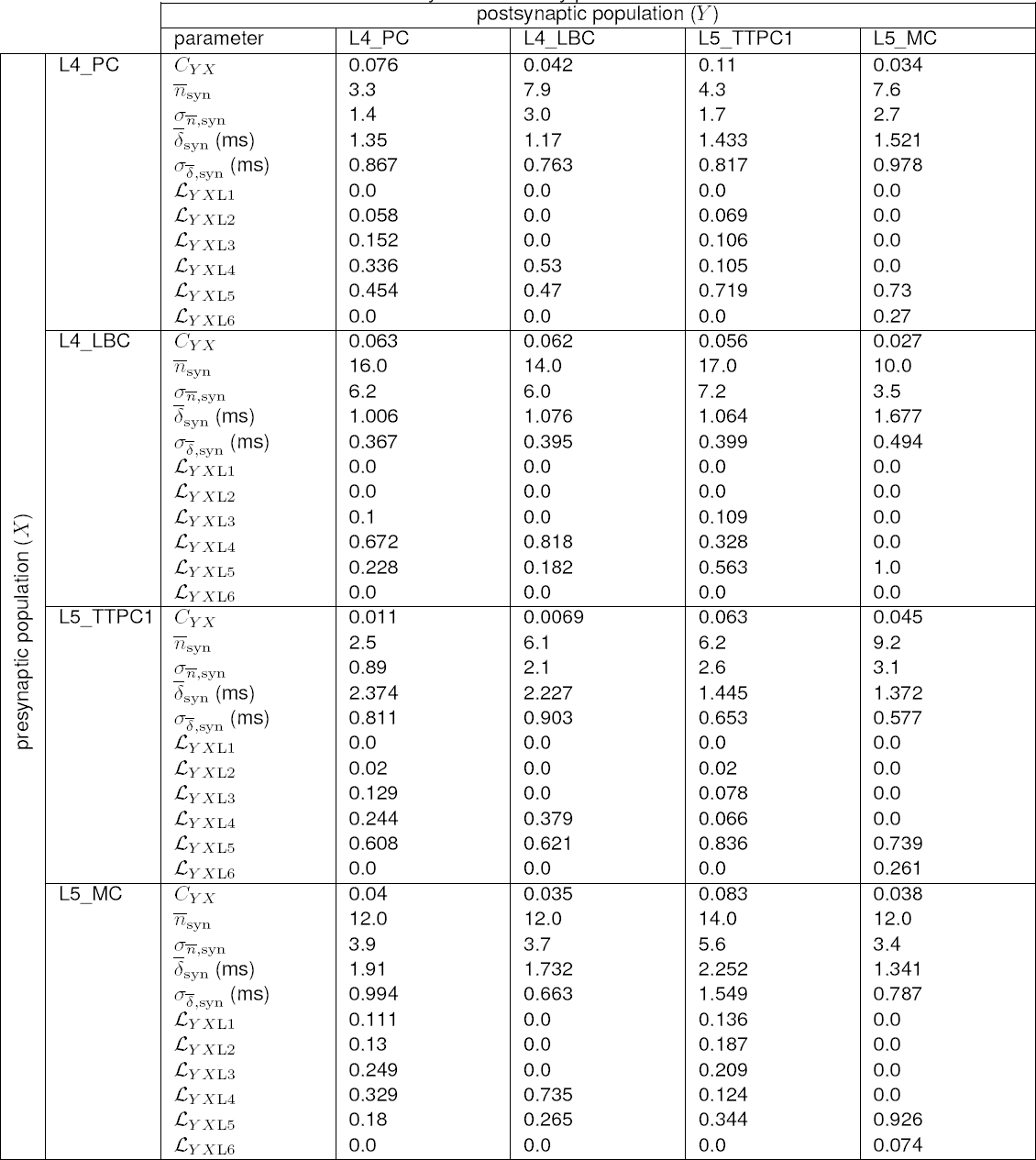
Summary of connectivity parameters.

##### Computation of extracellular potentials inside cortical column

For our multicompartment neuron network we chose to compute the extracellular potential vertically through the center of the column, with the most superficial contact at the top of layer 1 (*z* = 0) to a depth of *z* = −1500 *µ*m within layer 6. The inter-contact distance was Δ*_z_* = 100 *µ*m, and contacts were assumed to be circular with radius *r*_contact_ = 5 *µ*m and surface normal vectors aligned with the horizontal *y*-axis. For the electrode surface averaging we used *m* = 50 (cf. Eq (8) and Lindén et al. [2014]). For the calculation of extracellular potential inside the cortical column we assumed a homogeneous, isotropic, linear and ohmic extracellular conductivity *σ*_e_ = 0.3 S/m.

##### Computation of ECoG signal from Method-of-Images

The extracellular potential on top of cortex (ECoG) was computed by means of the Method-of-Images (MOI, see Section 2.2.2). In the example, the conductivity below the contact was set as *σ*_*G*_ = *σ*_*T*_ = 0.3 S/m, corresponding to the gray-matter value used above, while the conductivity on top of cortex was to set to be fully insulating, i.e., *σ*_*T*_ = 0 S/m. This could correspond to the situation where a grid of ECoG contacts are embedded in an insulating material (see, for example, Castagnola et al. [2014]). We further considered a single circular ECoG disk electrode with contact radius *r* = 250 *µ*m with its surface normal vector perpendicular to the brain surface. The disk electrode was centered at the vertical population axis and positioned at the upper boundary of layer 1. For the disk-electrode approximation (cf. Eq (8)) we set *m* = 500. (Note that the present MoI implementation requires all transmembrane currents to be represented as point sources confined within the boundaries of the middle (cortical) layer.

##### Computation of EEG and MEG signals

The most direct approach for computing EEG and MEG signals would be to (i) compute the per-neuron current dipole moment, (ii) compute the contribution to the signals from each neuron, and (iii) sum these signals to get the total EEG and MEG signals from the entire network. To reduce the computational demands, we instead compute the per-population current dipole moment **p***X* (*t*) using Eq (12). The total current dipole moment is then obtained by summing over all populations, i.e., **p** = Σ_*X*_ **p***X*.

From **p***X* we computed the EEG (surface electric potentials on the scalp layer) of the four-sphere head model as described above, and similarly magnetic fields **B**_**p**_. For the four-sphere head model we assumed conductivities *σ*_*s*_ *ϵ*{0.3, 1.5, 0.015, 0.3} S/m and radii *rs ϵ*{79, 80, 85, 90} mm for brain, cerebrospinal fluid (CSF), skull and scalp, respectively [Nunez and Srinivasan, 2006; Næss et al., 2017]. We positioned each population current dipole **p***X* below the brain-CSF boundary on the vertical *z*-axis (thus *x* = *y* = 0) at *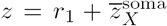*, where *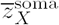* was the average soma depth within each population. Surface potentials, i.e., EEG potentials, and magnetic fields where computed for polar angles *θ ϵ* [ *π/*4, *π/*4] with angular resolution Δ*θ* = *π/*16 as illustrated in Fig 5E (azimuth angles φ = 0), resulting in a contact separation along the arc of *r*_4_*π/*16 ≈18 mm. Different magnetoelectroencephalogram (MEG) equipment may be sensitive to different components of the magnetic field [Hämäläinen et al., 1993]. We show different scalar components of the magnetic field computed on the surface of the four-sphere head model as described above (in Section 2.3.5).

**Figure 5:**
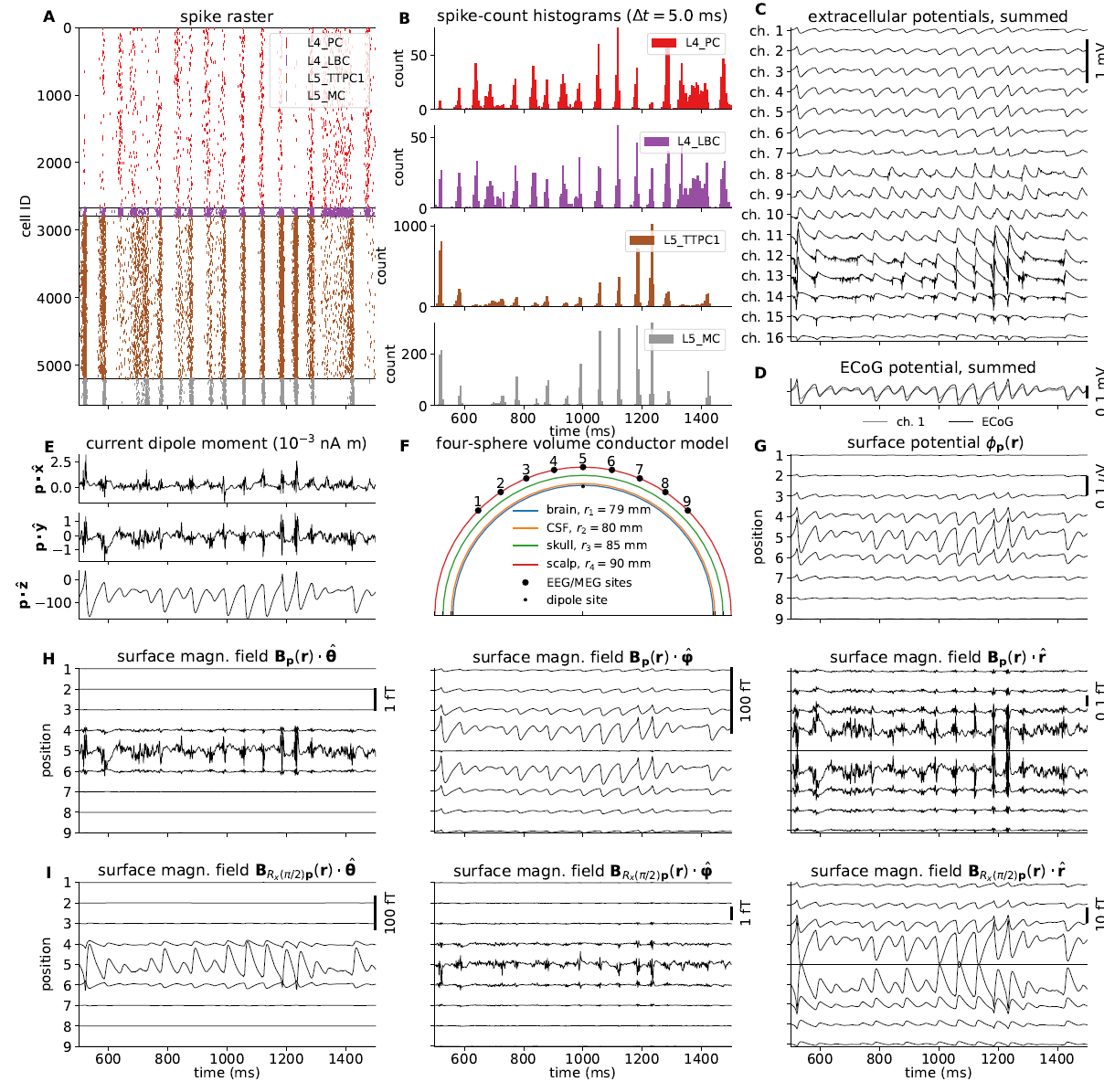
Intra-and extracellular measures of activity in example network. **A** Spike raster plot for each population. Each row of dots corresponds to the spike train of one neuron, color coded by population. **B** Population spike rates computed by summing number of spike events in each population in temporal bins of width Δ*t* = 5 ms. **C** Extracellular potentials as function of depth assuming an infinite volume conductor. **D** Extracellular potential on top of cortex (ECoG) assuming a discontinuous jump in conductivity between brain (*σ* = 0.3 S/m) and a non-conducting cover medium (*σ* = 0 S/m) and electrode surface radius *r* = 250 *µ*m. The signal is compared to the channel 1 extracellular potential in panel C (gray line). **E** Component-wise contributions to the total current dipole moment **p**(*t*) summed over population contributions. **F** Illustration of upper half of the four-sphere head model (with conductivities *σ*_*s*_ ϵ {0.3, 1.5, 0.015, 0.3}S/m and radii *r*_*s*_ ϵ {79, 80, 85, 90} mm for brain, csf, skull and scalp, respectively), dipole location in inner brain sphere and scalp measurement locations. The sites in the *xz*-plane numbered 1-9 mark the locations where electric potentials and magnetic fields are computed, each offset by an arc length of *r*_4_*π/*16 ≈18 mm. **G** EEG scalp potentials from multicompartment-neuron network activity with radially oriented populations. **H** Tangential and radial components of the head-surface magnetic field (MEG) from multicompartment-neuron network activity with radially oriented population. **I** Tangential and radial components of the magnetic field (MEG) on the head surface, with underlying dipole sources rotated by an angle *θ* = *π/*2 around the *x*-axis (thus with apical dendrites pointing into the plane). (Note that at position 5, the unit vectors 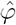 and 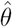 are defined to be directed in the positive *y*-and *x*-directions, respectively.)

##### Simulation details

Simulations were run for a total duration of *T* = 1500 ms with a simulation step size *dt* = 0.0625 ms (16 kHz sampling frequency). The first 500 ms were discarded as startup transient. All neurons were initialized at a membrane voltage *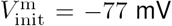* mV and temperature *T*_celsius_ = 34^*v*^C (affecting membrane-channel dynamics).

##### 2.7. Technical details

All source codes and development history of past and present versions of LFPy are publicly available on GitHub (see https://github.com/LFPy/LFPy), using ‘git’ (https://git-scm.com) for code provenance tracking. LFPy is released with an open-source software licence (GPL), which alongside GitHub functionality for listing issues, integration with automated testing, easy forking, local development and merges of upstream changes, facilitates continued, community-based LFPy development.

LFPy2.0 requires Python (continuously tested w. v2.7, v3.4-3.6), an MPI (message-parsing interface) implementation such as OpenMPI, NEURON v7.4 or newer compiled with MPI and bindings for Python, Cython, and the Python packages mpi4py, numpy, scipy, h5py, csa (https://github.com/INCF/csa) and Neu-roTools (http://neuralensemble.org/NeuroTools). In order to run all example files also matplotlib and Jupyter (http://jupyter.org) have to be installed, but prebuilt Python distributions such as Anaconda (https://www.continuum.io) should provide these common Python packages, or provide easy means of installing LFPy dependencies (issuing, for example, “conda install mpi4py” on the command line). Detailed instructions for installing dependencies for common operating systems (MacOS, Linux, Windows) are provided in the online LFPy documentation (http://lfpy.readthedocs.io).

The latest stable LFPy release on the Python Package Index (https://pypi.python.org) can be installed by issuing:

~~~
$ pip install LFPy --user
~~~

which may prompt the install of also other missing dependencies. The command

~~~
$ pip install --upgrade --no-deps LFPy
~~~

may be used to upgrade an already existing installation of LFPy (without updating other dependencies). In order to obtain all LFPy source codes and corresponding example files, we recommend users to checkout the LFPy source code on GitHub, after installing the git version control software:

~~~
$ **cd** <path to repository folder>
$ git clone https://github.com/LFPy/LFPy.git
$ **cd** LFPy
$ pip install -r requirements.txt
$ python setup.py develop –user
~~~

More detail is provided on http://lfpy.readthedocs.io.

###### Reproducibility

The simulated results and analysis presented here were made possible using Python 2.7.11 with the Intel(R) MPI Library v5.1.3, NEURON v7.5 (1472:078b74551227), Cython v0.23.4, LFPy (github.com/LFPy/LFPy, SHA:0d1509), mpi4py v2.0.0, numpy v1.10.4, scipy v0.17.0, h5py v2.6.0, parameters (github.com/NeuralEnsemble/parameters, SHA:v0aaeb), csa (github.com/INCF/csa, SHA:452a35) and matplotlib v2.1.0 running in parallel using 120-4800 cores on the JURECA cluster in Jülich, Germany, composed of two 2.5 GHz Intel Xeon E5-2680 v3 Haswell CPUs per node (2 × 12 cores), running the CentOS 7 Linux operating system. Each node had at least 128 GB of 2133 MHz DDR4 memory. All software packages were compiled using the GNU Compiler Collection (GCC) v4.9.3. All source codes for this study will be provided as LFPy example files on GitHub.

## 3. Results

### 3.1. Single-neuron activity and extracellular measurements

The first version of LFPy [Lindén et al., 2014] assumed the active model neurons to be embedded in an infinite homogeneous volume conductor and was most suited to compute extracellular potentials (spikes, LFPs) inside the brain. One new feature of LFPy2.0 compared to the first version of LFPy is that electrical potentials outside cortex (ECoG, EEG), as well as magnetic fields both inside and outside cortex (MEG), can be computed. These new measures are illustrated in Fig 2 for a single synaptically activated pyramidal neuron.

Fig 2A presents a basic LFPy simulation example where a passive neuron model with simplified morphology receives a single synaptic input current (inset I). We computed the extracellular potential in the *xz*-plane (color image plot), using the assumption of line sources for each dendritic compartment, a spherical current source representing the soma, and homogeneous conductivity (Eq (7)). The postsynaptic response is reflected as a somatic depolarization (inset II) and as a deflection in the extracellular potential in the location **r** (blue dot, inset III). The corresponding current dipole moment **p**(**r***, t*) was computed using Eq (12) and is illustrated by the black arrow. The *x*- and *z*-components of the current dipole moment are illustrated in inset IV, and we note the much larger dipole moment component in the vertical *z*-direction compared to the lateral *x*-direction. We do not show the *y*-component of the current dipole moment as all segments in this simplified neuronal morphology are located in the *xz*-plane (hence 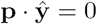.

To illustrate the fact that a current dipole potential (Eq (15)) gives a good approximation to the extracellular potential *φ* far away from the neuron, we compare with results from using the more comprehensive line-source method (Eq (6)) in panel B: The line-source potential *φ* is shown inside the dashed circle of radius *r >* 500 *µ*m, while the dipole potential *φ*_**p**_ is shown outside the circle. The inset shows the dipole potential corresponding to the colored dots located at a distance of 750 *µ*m.

In panel C we similarly compute the magnetic field for radii *r >* 500 *µ*m using the current dipole moment (Eq (30)), and axial currents inside (Eq (32)). The axial currents were computed from per-compartment membrane potentials as described in Section 2.1.2. For both color image plot and the inset, we show the dominating magnetic field component, i.e., the *y*-component. As for the electrical potential in panel B, we see that the predicted magnetic fields match well at the *r* = 500 *µ*m interface.

Panel D illustrates the layout of scalp-layer measurement sites on the four-sphere head model described in Section 2.3.3. The numbered points along the outer scalp layer represents measurement locations for EEG and MEG signals. The single current dipole moment is positioned beneath the CSF-brain boundary on the vertical *z*-axis (see caption for details). Panel E shows the corresponding scalp surface potentials which is dominated by the *z*-component of the current dipole moment (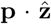, panel A inset IV). Panel F shows the corresponding dominant tangential magnetic field component (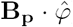) computed from the current dipole moment using Eq (30). At the center location (location 5) only the *x*-component (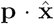) contributes to the signal, in the other locations both the *x*-and *y*-components contribute.

### 3.2 Network activity and extracellular measurements

The second main new feature of LFPy2.0 is the possibility to simulate recurrently connected networks of neurons in parallel. Our exemplary network, shown in Fig 4, demonstrating this new feature is based on a subset of cortical single-cell models, synapse models and connectivity data from Markram et al. [2015] obtained from The Neocortical Microcircuit Collaboration (NMC) Portal [Ramaswamy et al., 2015]. The implementation is described in detail in Section 2.6.

In addition to supporting simulations of neuronal networks with biophysically detailed single-neuron models in parallel, LFPy2.0 allows for concurrent calculations of extracellular measures of network activity. Specifically, the extracellular potentials at specific positions can be computed at each time step which avoids the memory-demanding process of recording transmembrane currents in all compartments for the duration of the simulation, either to disk or to memory. In the present example, the current dipole moment was calculated at every time step, and this amounted to a useful dimensionality reduction, as only the *x, y, z*-axis components of **p** per population *X* had to be stored. Assuming serial execution, then for each neuron population *X*, the total memory consumption is then reduced by a factor 3*/*(*N*_*X*_ *n*^seg^) where *N*_*X*_ is the population size and *n*^seg^ the number of compartments per neuron (see panel A for values), compared to storing currents. The per-population current dipole moments were then used to predict EEG scalp surface potentials and MEG signals in the corresponding locations. Note that per-population current dipole moments can be stored, EEG and MEG signal can be computed with other head models at a later stage.

#### 3.2.1. Network spiking activity

Fig 5 shows the various predicted measurements for a one-second period of network activity. The spike raster and corresponding spike-count histogram (panels A and B) demonstrate the network’s tendency to produce synchronous irregular patterns of activity with the parameterization summarized in Section 2.6, Tables 1-3 and Fig 4B. The per-neuron spike occurrences in the excitatory populations L4_PC and L5_TTPC1 were sparser than for the inhibitory populations L4_LBC and L5_MC. As in the full circuit of Markram et al. [2015], it is possible that an asynchronous network state could have been obtained by modifying extracellular [Ca2+]o-dependent release probabilities *P*_*u*_ for the different synapse types in the model [Borst, 2010; Markram et al., 2015]. A modification of release probabilities can shift the effective balance between excitatory and inhibitory synapse activations, but also incorporation of a larger sample of heterogeneous cell types in the model could have brought the network into an asynchronous state, essentially by increasing the amount of inhibitory feed-back. In particular interneuron expression in neocortex is known to be more heterogeneous and more dense than demonstrated here [Markram et al., 2004, 2015]. However, as our main focus here is to present new simulation technology now incorporated in LFPy, we did not pursue this line of inquiry.

#### 3.2.2. Local field potentials (LFPs)

The extracellular potentials as would be measured by a 16-channel laminar probe positioned through the center axis of the cylindrical column, are shown in Fig 5C. The computed extracellular potentials are observed to be of the same order of magnitude as experimentally measured spontaneous potentials (≃ 0.1– 1 mV, see Maier et al. [2010]; Hagen et al. [2015]; Reyes-Puerta et al. [2016]). We further observe that the synchronous events seen in the spiking activity (panel A) are reflected as substantial fluctuations in the extracellular potential with amplitudes close to 0.5 mV.

The signals in neighboring channels are further observed to be fairly correlated with comparable amplitudes, irrespective of the presence of somatic compartments at the depths of the contacts (Fig 4F). At the superficial channels 1–6, deflections in the electric potential following synchronous network activation are predominantly negative, while a change in sign occur around channel 7 (near the boundary between layer 3 and 4). The strongest deflections of the extracellular potential are typically observed at contacts within layer 5 (ch. 11–13), that is, at depths corresponding to the dense branching of basal dendrites and somas of the large layer 5 pyramidal neuron population. These deflections reflect that the soma compartments and basal dendrites are expected to act as dominant sources of the transmembrane currents setting up the extracellular potential [Lindén et al., 2010]. Adding further to this, layers 4 and 5 also had the highest overall densities of excitatory and inhibitory synapses in the present model (Fig 4G). Some spike events (extracellular signatures of action potentials) are seen in ch. 14, produced by one or several neurons located near the virtual recording device.

Further investigation of the different contributors (Fig 6A-D) to the extracellular potential (Fig 5C), revealed that most of the signal variance across depth can be explained by transmembrane currents of the two excitatory populations (Fig 6E). Even if the cell numbers in the two pyramidal-cell population were similar, population L5_TTPC1 contributed more to the signal than population L4_PC at all channels except at channel 9 (around which the L4_PC somas are positioned).

**Figure 6:**
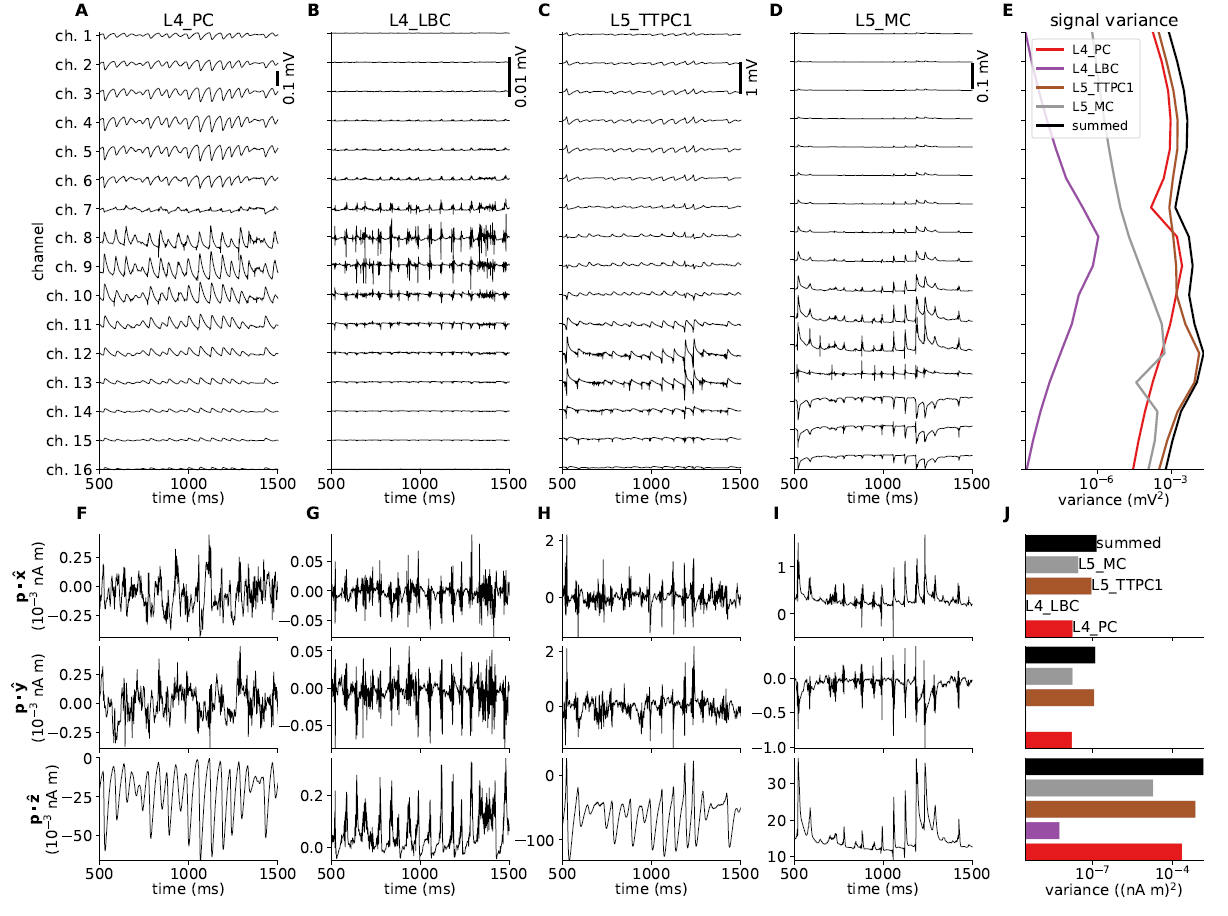
Per-population contributions to the extracellular potential and current dipole moment and corresponding signal vari-ance. **A-D** Contributions to the extracellular potential from populations *X* ϵ{L4_PC, L4_LBC, L5_TTPC1, L5_MC} in the network across depth. **E** Extracellular potential variance across depth for contributions of each population, and for the sum over populations. **F-I** *x, y, z*-components of the per-population contribution to the summed current dipole moment. **J** Per-component current dipole moment variance for each population and for summed signals.

#### 3.2.3. ECoG signal

Fig 5D compares the extracellular potential in the topmost channel 1 (gray line), predicted under the assumptions of dendritic line sources, somatic spherical sources and an infinite homogeneous extracellular medium (cf. Eq (7)), with our ECoG prediction at the same depth (black line). The ECoG signal was computed assuming a wide contact (*r*_contact_ = 250 *µ*m) aligned horizontally on top of a flat cortex (*z* = 0). Further, for the ECoG signal the method-of-images (MoI; cf. Eq (11)) was used to account for a conductivity discontinuity at the cortical surface. Here, zero conductivity (mimicking, for example, the situation with an insulating mat surrounding the ECoG contact [Castagnola et al., 2014]) was assumed above the cortical surface, while the grey-matter value of *σ*_e_ = 0.3 S/m was assumed below.

The amplitude of the ECoG trace was slightly increased compared to the potential measured by the smaller electrode. This amplitude increase can be attributed to the fact that a reduction in conductivity above the boundary would decrease the value of the denominator of Eq (11), and hence increase the signal amplitude below insulating cortical surfaces [Pettersen et al., 2006]. The expected increased signal amplitude from this conductivity step is here counter-measured by the larger diameter of the ECoG contact (*r*_contact_ = 250 *µ*m vs. *r*_contact_ = 5 *µ*m) resulting in an increased average distance from the signal source to the contact point averaged over the contact’s surface. Detailed investigation of each signal normalized to the same standard deviation (not shown) revealed virtually indistinguishable features across time and in their power spectra.

#### 3.2.4. Current dipole moments

Fig 5E shows the three components of the total current dipole moment **p** stemming from the network activity. The most striking feature is the much larger *z*-component compared to the lateral *x*-and *y*-components. This large difference in component size, about two orders for magnitude, reflects (i) that the vertically aligned pyramidal cell morphologies span across several layers, and (ii) the near rotational symmetry of the model populations around the *z*-axis. Unlike the *z*-component, the lateral components largely cancel out. In the same way as for the extracellular potential, the two pyramidal populations are also the dominant sources of the total dipole moment (Fig 6F-J). We also note that the *z*-component of the population current dipole moment generally dominates the other components of the population dipoles, with the exception of the L4_LBC population. Here all components are tiny, reflecting the stellate dendritic morphology and the evenly distributed synapses onto the neurons in this population.

For our model network we note that the maximum magnitude of the current dipole moment is about 0.1 nAm, which is about two orders of magnitude smaller than previously estimated typical ‘mesoscopic’ dipole strengths [Hämäläinen et al., 1993, p. 418].

#### 3.2.5. EEG signals

As a demonstration of predicting non-invasive electric (‘EEG’) signals outside of the brain with LFPy2.0, we utilized the four-sphere head model (as implemented in class `FourSphereVolumeConductor`, see Methods) and defined scalp-layer measurement locations as illustrated in Fig 5F. We assumed the modeled network to represent a piece of cortical network positioned at the top of a cortical gyrus, so that the population axes were in the radial direction of the spherical head model. The current dipoles (computed above) were positioned below the interface between the CSF and the brain, more specifically the layer-4 and layer-5 population dipoles were positioned at the depth of the center of layer 4 and layer 5, respectively.

As observed in Fig 5G, the temporal form of the scalp potentials corresponds directly to the temporal form of the dominant *z*-component of the current dipole moment in panel E. For an infinite volume conductor it follows directly from Eq (15) that the recorded scalp potential will be proportional to this dipole moment at recording positions directly (radially) above the dipole location. Likewise, inspection of the formulas for the four-sphere head model shows that this is also the case for the scalp-potential contributions from both the radial (Eq (16)–Eq (17)) and tangential (Eq (18)–Eq (19)) dipole components (although with different proportionality constants for the two components).

For the present example network comprising 5594 neurons of which 5077 are pyramidal cells, we observe the magnitudes of the fluctuating scalp potential directly on top of the dipole sites to be on the order of 0.1 *µ*V. This is about two orders of magnitude smaller than the typical size of measured EEG signals of 10 ∼*µ*V [Nunez and Srinivasan, 2006, Fig. 1.1].

The weakly conducting skull layer (compared to the highly conductive brain, spinal fluid and scalp layers) results in a spatial ‘low-pass filter effect’ from volume conduction [Nunez and Srinivasan, 2006, Ch. 6]. This low-pass effect accounts for the relatively weak attenuation of the EEG signal with lateral distance from the center position (position 5 in panel F) along the head surface, as observed in panel G. On the surface of a spherical volume conductor with homogeneous conductivity inside the sphere, but otherwise zero conductivity outside the sphere’s surface (1-sphere head model), the potential from a current dipole would decay in amplitude at a higher rate compared to our 4-sphere head-model case with a spherical skull layer with low conductivity. However, in an infinite homogeneous volume conductor the decay in electric potential along the putative sphere’s surface would decay with a lower rate than both the 1-sphere and 4-sphere head models, see Nunez and Srinivasan [2006, Ch. 6] for a comparison.

#### 3.2.6. MEG signals

The computed current dipole moments in Fig 5E was also used to compute MEG signals. Panel H shows the computed magnetic fields for the same set-up providing the EEG signals in panel G, that is, radially oriented population current dipoles. In this situation the only sizable magnetic field is directed in the tangential direction around the vertical *z*-axis. With our spherical coordinates this corresponds to the *φ*-direction where the unit vector 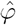 points in counter-clockwise direction. Note also that the magnetic field is almost zero straight above the dipole (position 5), as here the vectors **p** and **R** are near parallel so that the vector product in Eq (30) is very small. We also observe that the magnetic field is symmetric around the center position (position 5), so that the field at position 6 is always similar to the field at position 4, and so on.

For EEG signals, equivalent radial dipoles located at the ‘crowns’ of gyri are generally expected to give the largest signal contributions [Nunez and Srinivasan, 2006]. For MEG signals, on the other hand, equivalent current dipoles in brain sulci oriented tangentially to the head surface is expected to provide the largest signals [Hämäläinen et al., 1993]. In Fig 5I we thus show the magnetic field with the current dipole moment in panel E directed in a tangential direction (that is, in the *y*-direction into the paper in panel F) rather than in the radial direction. In this situation the largest magnetic field component is in the tangential direction 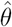 (around the *y*-axis) in position 5. The 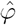-component is as expected negligible, while the radial component is antisymmetric around position 5, but negligible in position 5.

Typical magnetic fields measured in human MEG are on the order of 50–500 fT [Hämäläinen et al., 1993], and in Fig 5I we find that magnetic fields of similar magnitudes (∼ 100 fT) are predicted when the current dipole moment from our network is oriented in parallel to the cortical surface. Note, however, that in our model set-up, the dipole is only 11 mm away from the closest MEG sensor at position 5, while in human recordings the minimum distance between tangential dipoles in brain sulci and the MEG sensors may be several centimeters [Hämäläinen et al., 1993]. As the magnetic field from a current dipole decays as the square of the distance (see Eq (30)), our model likely gives an overestimate of the contribution to the MEG signal from our model network when applied to a human setting.

In Fig 5H we also observe sizable magnetic fields (∼ 20–40 fT) generated by radially-oriented current dipoles. However, the generated fields are in the angular *φ*-direction where the fields have opposite directions on each side of the central position (position 5). Thus, in a setting with several such neighbouring dipoles (generated by neighbouring populations) on cortical gyri, there will be large cancellations effects. Despite the larger distances from the MEG sensors, tangentially oriented dipoles in sulci is therefore expected to dominate the measured MEG in human settings [Hämäläinen et al., 1993].

### 3.3. LFPy parallel network performance

In order to assess the performance figures of multicompartment-neuron network implementations in LFPy on a high-performance computing (HPC) facility, we performed a series of simulations with two-population versions of the network presented above. These modified networks consisted only of the layer-5 m-types L5_TTPC1 and L5_MC. We modified cell counts per population *N*_*X*_ and connection probabilities *C*_*Y X*_ de-pending on chosen network population sizes *N*_*X*_ as noted in the text below. All other simulation parameters were kept fixed as given in Tables 1-3.

First, we compared set-up times, creation times of populations and connections, and simulation times for instantiations of similarly sized reference networks 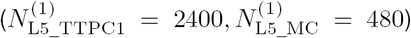 for different number of MPI processes *N*_MPI_ (Fig 7A). *N*_MPI_ was set identical to the number of available physical cores (no multi-threading). A seed value for the random number generator for each network instantiation was varied to obtain an *N* = 3 sample size for each tested value of *N*_MPI_. Both with predictions of extracellular potentials and current dipole moments (continuous lines) and without (dotted lines), the biggest fraction of the total computational time was spent during the main simulation part (red curves), that is, where the simulation is advanced time step by time step. The additional computational cost of computing extracellular potentials and current dipole moments was less than half compared to just simulating the spiking activity in the recurrently connected network. The times spent creating all recurrent connections and synapses (green curves) were between a factor 16 and 32 shorter than the simulation time.

**Figure 7:**
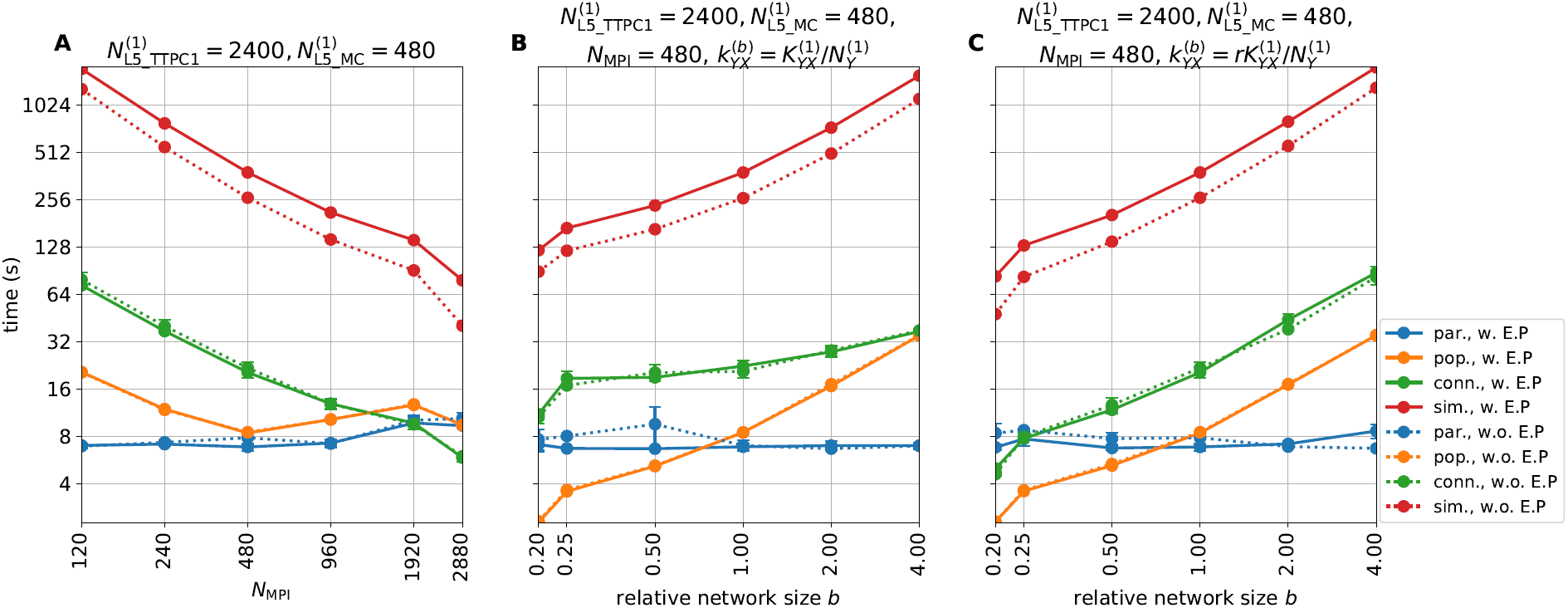
Parallel performance with networks in LFPy. **A** Initialization of parameters (par.), population create (pop.), connectivity build (conn.) and main simulation time (sim.) as functions of number of physical CPU cores/MPI processes (*N*_MPI_). The reference network population sizes *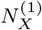* for {*X* ϵ {L5_TTPC1, L5_MC} are given in the panel title. The network was otherwise constructed with synapse, stimulus and connectivity parameters for each possible connection as given in Tables 1-3. Times shown with continuous lines were obtained for simulations that included calculations of extracellular potentials and current dipole moments as in Figures 2-6 (w. E.P.), while times shown with dotted lines were obtained for simulations with no such signal predictions (w.o. E.P.). Each data value is shown as the mean and standard deviation of times obtained from *N* = 3 network realizations instantiated with different random seeds. **B** Initialization of parameters, population create, connectivity build and main simulation time as functions of network size relative to the reference network population sizes *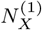* for *X* ϵ{L5_TTPC1, L5_MC} as given in the panel title. The superset ‘(1)’ denotes a relative network size *b* = 1. Simulations were run using a fixed MPI process count *N*_MPI_ and connection probabilities *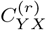* were recomputed for different values of *b*, such that the expected total number of connections *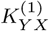* was constant between each simulation (using Eq (34)). The set-up was otherwise identical to the set-up in panel A. **C** Same as panel B, but with a fixed expected per-cell synapse in-degree 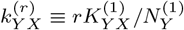 across different relative network sizes.

The creation of connections and simulation times scaled strongly with *N*_MPI_. An optimal, or strong, log-log-linear scaling curve can be represented as a function *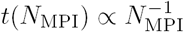*, in particular for *N*_MPI_ ≤480, as these *N*_MPI_-values result in an even load balance across parallel processes with the presently used round-robin distribution of cells across MPI processes (see Section 2.5 for details). Each parallel process has the same number of cells of each m-type, segments 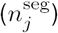 and state variables corresponding to different active ion-channel models. Only variations in per-cell in-degrees (synapse counts) across different processes and simulations occurred due to the random network connectivity model, but even with different random seeds in each trial the trial variability was small (error bars denoting standard deviations are hardly seen).

The creation of populations (orange curves) however showed worse scaling behaviour for *N*_MPI_ > 480, in part due to uneven load balance. Another possible reason for reduced performance was the increased strain on the file system as all processes simultaneously access the same single-neuron source files upon instantiating individual `NetworkCell` objects. This might have been avoided by creating local copies of the necessary files on each compute node, but we did not pursue this here as the overall time spent instantiating neuron populations was only a fraction of the observed simulation times. The loading of parameters and other needed data (blue curves) was, as expected, fairly constant for different values of *N*_MPI_ as we did not parallelize the corresponding code.

As a second scaling-performance test, we ran series of simulations with *N*_MPI_ = 480 but varied the total network size by a factor *b* ϵ{0.2, 0.25, 0.5, 1, 2, 4} while keeping the expected number of connections *K*_*Y X*_ (and thus the number of synapses) between pre-and post-synaptic populations *X* and *Y* fixed (Fig 7B). The expected number of randomly created (binomially distributed) connections *K*_*Y X*_ was calculated using the relation [Potjans and Diesmann, 2014]:

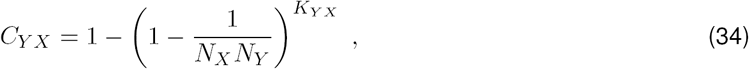

with reference network size 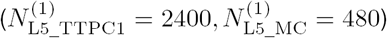 and connection probabilities *C*_*Y X*_ as givenin Table 3. Similar to the test presented in panel A, most of the total computation time was spent during the main simulation part (red curves), followed by creation of connections (green curves) and loading of different parameters (blue curves).

In contrast to the previous case, the creation of cells in the network displayed strong scaling with network size (which implies a relationship *t*(*r*) ∝*b*). The supra-optimal scaling seen for connections can be explained by the creation of similar connection counts across different factors *b*. (Note that supra-optimal scaling implies that *t*(*r*) ∝*b*^*q*^ with exponent *q* ϵ (0, 1), while sub-optimal scaling implies that *q >* 1.) For the tested factors *b* = 0.25 and *b* = 0.5 we expected sub-optimal scaling for creating populations and connections, as well as for simulation duration. These *b*-values gave different cell counts and thus inhomogeneous load-balances across MPI processes, which was unavoidable with the presently used round-robin parallelization scheme. A jump in performance was seen for *b* = 0.2 which resulted in only one multicompartment neuron and corresponding calculations on each MPI process.

As a third scaling-performance test we fixed the mean per-cell synapse in-degree *k*_*Y X*_ (count of incoming connections per cell) and reran network simulations for different network sizes (Fig 7C). The total number of connections was thus set to 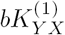 and corresponding connection probabilities *C*_*Y X*_ were recomputed accordingly using Eq (34). As expected, this modification mostly affected the time spent creating connections (green curve), and resulted in a near-linear performance curve for scaling factors *b* ≥ 1.

As a final performance assessment we repeated the experiment described above with upscaled networks and increased MPI pool sizes. In Fig 8A we set the reference network population sizes *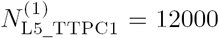* and *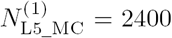* and varied *N*_MPI_ between 600 and 4800. LFPy’s parallel performance was strong also here, and Fig 7A consequently shows trends similar to the findings for the smaller network. Here, the time spent creating populations (orange curves) was reasonably invariant for different *N*_MPI_ values, and increased overall by some factor 2-4 compared to the previous case. The parameter loading times were similar, while the time spent connecting the network was increased by a factor ∼4, but the simulation times increased only by a factor ≲ 2. The differences in connection and simulation times seen here, can be explained by the fact that the typical synapse in-degrees were not preserved. Instead, the synapse in-degrees were increased for the larger network, as we used the connection probability values defined in Table 3.

**Figure 8:**
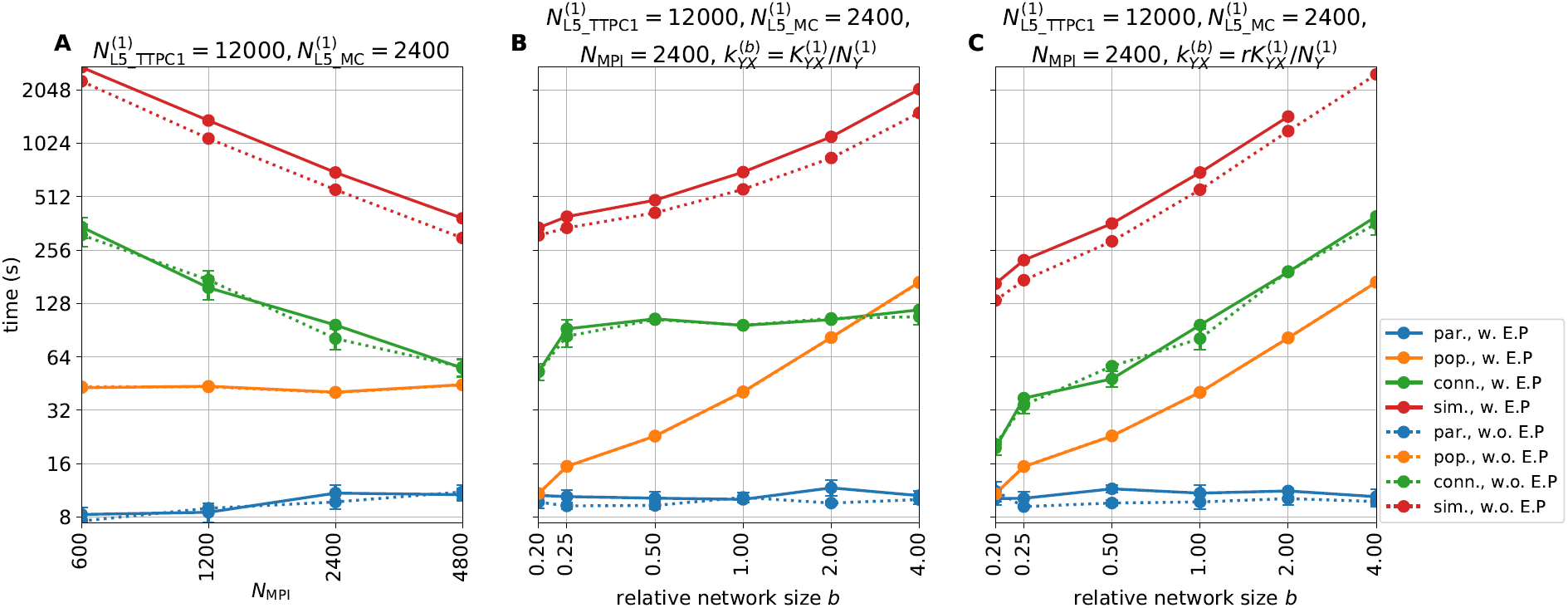
Parallel performance with networks in LFPy II. **A** Similar to Fig 7A, but with network population sizes upscaled by a factor 5, and a corresponding increase in parallel job sizes. **B-C** Similar to Fig 7B-C, but with network population sizes and parallel job sizes increased by a factor 5.

In Fig 8 panels B and C we set *N*_MPI_ = 2400, and varied the network population sizes relative to the reference network population sizes in panel A by the factor *b* ϵ{0.2, 0.25, 0.5, 1, 2, 4}. Again, the performance figures were in qualitative agreement with the previous results for the smaller network and smaller MPI pool sizes. The population creation times and simulation times with and without signal predictions displayed strong scaling with relative network size. The time spent loading parameters was increased by a small amount (by a factor ≲ 2), which likely reflected the increased strain on the file and communication system on the cluster, due to larger MPI pool sizes. The times spent creating the populations were also here near ideally dependent on *N*_MPI_ in both panels B and C. As the total number of connections (and synapses) were conserved across network population sizes in panel B, the connection times varied only by a factor two from the smallest to the largest network. In panel C, where the number of connections per neuron was kept approximately constant, a doubling in network size resulted in a doubling in connection times. The larger network simulations required approximately twice the amount of time, compared to the smaller network simulations in Fig 7. In panel C, simulations with LFP predictions consistently failed for the largest network size (*b* = 4), most likely due to lack of available memory to create arrays for storing current dipole moments and extracellular potentials with the increased count of instantiated connections.

## 4. Discussion

In the present paper we have presented LFPy2.0, a majorly revised version of the LFPy Python package with several added features compared to its initial release [Lindén et al., 2014].

### 4.1. New features in LFPy2.0

The first version of LFPy only allowed for the computation of electrical measurements from activity in single neurons or, by trivial parallellization, populations of neurons only receiving feedforward synaptic input. LFPy2.0 allows for simulations of recurrently connected neurons as well, for example the types of neuronal networks in cortex. Further, the first version of LFPy was tailored to compute extracellular potentials (spikes, LFPs) inside the brain. Here it was assumed that all active neurons were embedded in an infinite *homogeneous* (i.e., same extracellular conductivity everywhere) and *isotropic* (i.e., same extracellular conductivity in all directions) volume conductor (Section 2.2.1). LFPy2.0 includes several new features and measures of neural activity:

- Stepwise discontinuities in the extracellular conductivity, such as at the cortical surface, can be included by means of the Method-of-Images (Section 2.2.2) to compute potentials immediately below or on the cortical surface (i.e., electrocorticographic recordings; ECoG). This approach can also be applied in the computation of potentials recorded by microelectrode arrays (MEAs) [Ness et al., 2015].
- Cylindrical anisotropic conductivity (Section 2.2.3) can be included in the computation of spikes and LFPs, reflecting for example that in cortex and hippocampus the conductivity might be larger in the depth direction (along the apical pyramidal-neuron dendrites) than in the lateral directions [Goto et al., 2010].
- Current dipole moments from single neurons and populations of neurons are computed (Section 2.3.1) for later use in calculation of signals of systems-level electrical and magnetic recordings (EEG, ECoG, MEG), also for more detailed head models than what is considered presently in LFPy2.0 (as described in next two items).
- Electrical potentials at the scalp (electroencephalographic recordings; EEG) are computed from the current dipole moments and spherical head models, in particular the four-sphere head model [Nunez and Srinivasan, 2006; Næss et al., 2017], cf. Section 2.3.3. This four-sphere head also predicts ECoG signals (Section 2.3.4).
- Magnetic fields outside the head (magnetoencephalographic recordings; MEG) can be computed from the current dipole moments assuming a spherically symmetric head model (Section 2.3.5). Likewise, magnetic field inside the brain can be computed directly from neuronal axial currents (Section 2.4).

LFPy2.0 also includes much more rigorous code testing with more than 260 unit tests, automated build testing with TravisCI (travis-ci.org/LFPy/LFPy) with different versions of Python (2.7, 3.4-3.6), test coverage of code using coveralls (coveralls.io/github/LFPy/LFPy), automated documentation builds using Read the Docs (http://lfpy.readthedocs.io), and several updated example files, as well as new examples demonstrating different scientific cases using the new functionalities. The software runs on a wide variety of operating systems, including Linux, Mac OS and Windows.

### 4.2. Example applications

To illustrate some of the new measurement modalities incorporated in LFPy2.0 we showed in Fig 2 the LFP and EEG signature of a simple pyramidal-like neuron receiving a single excitatory synaptic input on its apical dendrite. In this example the extracellular medium was assumed to be homogeneous, and a characteristic dipolar profile was observed in the extracellular potential (panel B). The accuracy of the far-field electrical dipole approximation (Eq (15)) for distances of a few millimeters or more away from the neuronal source, was also demonstrated. The corresponding magnetic field set up by the neuron (panel C) was quite distinct from the electric potential pattern, but also here far-field magnetic dipole approximation (Eq (30)) was observed to be accurate some distance away.

To illustrate the implementation of networks in LFPy2.0 we showed in Section 2.5 a code example for a small network using simplified ball-and-stick neurons connected by conductance-based synapses. Our main example applications were on a network of about 5500 morphologically and biophysically detailed neuron models from the reconstructed somatosensory cortex column of Markram et al. [2015], connected using probabilistic synapse models with short-term plasticity. For this example, Fig 6 provided results for a one-second epoch of network activity where spikes (panels A, B), LFPs inside the cortical model column (panel C), the ECoG signal recorded at cortical surface (panel D), and the net current dipole moment (panel E) were depicted. The computed current dipole moment was further used to compute the corresponding EEG signal with the four-sphere head model for the situation where the model network was placed on top of a cortical gyrus where the apical dendrites of the pyramidal neurons, and thus the current dipole moment, is pointing in the radial direction (panel G). The same current dipole moment was also used to compute the MEG signal, assuming a spherically-symmetric head volume-conductor model, both for the case when the net current dipole is directed perpendicular (panel H) and parallel (panel I) to the scalp. The latter situation could correspond to the case where the model network is positioned in a cortical sulcus.

While the example network was set up mainly to demonstrate the new features in LFPy2.0, some of the example results are notable. As expected the two excitatory pyramidal cell populations in the network provided almost all of the recorded LFP signal (except in the deep layers where the layer-5 inhibitory Martinotti-cell population also gave a sizable contribution), cf. Fig 7E. Likewise, the two excitatory pyramidal cell populations also gave the dominant contributions to the net current dipole moment providing the EEG and MEG signals (Fig 7J).

For the present example network comprising about 5000 pyramidal neurons, we observed the maximum magnitude of the EEG signal to be about 0.1 *µ*V (Fig 6G), that is, about two orders of magnitude smaller than the typical size of measured EEG signals of ∼10 *µ*V [Nunez and Srinivasan, 2006, Fig. 1.1]. Thus our example model network appears too small, that is, it incorporates too few pyramidal neurons, to account for the typical experimentally recorded EEG signal amplitudes.

The maximum magnetic field computed at the cortical surface was seen in Fig 5H–I to be about 100 fT, that is, similar in magnitude to typical magnetic fields measured by MEG sensors in a human setting (∼ 50– 500 fT [Hämäläinen et al., 1993]). However, our model predictions assumed the minimum distance between the current dipoles and the magnetic-field recording device to be only about a centimeter, likely much smaller than the typical minimal distance between the dominant tangential dipoles in cortical sulci and the human MEG sensors. Since the magnetic field around a current dipole decays as the square of the distance, our modeling likely substantially overestimates the magnetic field that would produced by the computed current dipoles in a human setting.

### 4.3. Use of LFPy

#### Comparison of candidate models with experiments

An obvious application of LFPy is, following the tradition of physics, to (i) compute predictions of the various available measures of neural activity from different candidate models and (ii) identify which model, or which class of models, is in best agreement with the experimental data. While not always possible, the approach is preferably pursued on multimodal data measured simultaneously (for example simultaneous recordings of spikes, LFP and ECoG). The multi-objective comparison of experimental data with candidate models is a subject on its own, and will not be discussed here (but see, for example, Druckmann et al. [2007]).

#### Validation of data analysis methods

Neuroscience relies on data analysis, and data analysis methods should be validated [Denker et al., 2012]. An important application of LFPy could be to provide model-based ground-truth benchmarking data for such validation. This approach has already been used with biophysically detailed neuron models to test methods for spike sorting [Einevoll et al., 2012; Hagen et al., 2015; Lee et al., 2017], neuron classification [Buccino et al., 2017], estimation of firing rates from multi-unit activity (MUA) [Pettersen et al., 2008], current-source density (CSD) analysis [Pettersen et al., 2008; Łęski et al., 2011; Ness et al., 2015], independent component analysis (ICA) [Głąbska et al., 2014] and laminar population analysis (LPA) [Głąbska et al., 2016].

Likewise, LFPy could be used to aid in the interpretation of various statistical measures of electrophysiological activity such as *spike-triggered LFP* or *mutual information* [Einevoll et al., 2013]. The interpretation of these measures in terms of the underlying neural network activity is a priori not trivial, but intuition and understanding can be gained by LFPy model investigations where simulation results can be compared with neural activity directly. An example of this was given in Hagen et al. [2016]. There the spike-triggered LFP as measured in the model simulation was compared with other ways of accounting for spike-LFP relationships with a simpler physical explanation, that is, the LFP signature following activation of a presynaptic neural population.

It should be noted that the LFPy network model does not necessarily have to be finely tuned to a particular experimental system in order for it to be suitable for validation of data analysis methods: Methods claimed to have fairly general applicability should also be applicable to biologically plausible example network models.

#### Testing of simplified modeling schemes

LFPy now allows for the concurrent simulation of intracellular (membrane potential) and extracellular signals (spikes, MUA, LFP, EEG, MEG) for recurrent networks of biophysically and morphologically detailed neuron models. Such network models are computationally demanding to run [Markram et al., 2015], in particular when extracellular signals are computed simultaneously [Reimann et al., 2013]. A computationally less demanding alternative is a *hybrid LFP* scheme where the network dynamics, that is, spikes, are modeled with simple point-neuron models such as the integrate-and fire model, and the stored spikes are played back in a second computational step computing the extracellular potentials using multicompartment neuron models [Mazzoni et al., 2015; Hagen et al., 2016].

This scheme requires that salient features of spiking activity of networks of detailed multicompartment neuron models can be accurately captured by point-neuron network models. This was for example demonstrated by Rössert et al. [2016] who reproduced key network behaviour of a reconstructed somatosensory column [Markram et al., 2015] by systematic mapping of synaptic input to somatic responses in generalized leaky integrate-and-fire neurons. Likewise, the accuracy of the second step in the hybrid scheme where the extracellular potential is computed, can be systematically tested by comparing resulting predicted extracellular potential with the ground-truth potentials provided by LFPy. The same approach can naturally also be applied to test other simplified schemes for computing extracellular signals.

### 4.4. Possible refinements of measurement models in LFPy

#### Frequency-dependence of extracellular conductivity

The present forward-modeling schemes for electrical potentials assume the extracellular conductivities *σ*_e_ to be independent of frequency. If such a frequency dependence is found and described, it can in principle be straightforwardly incorporated by considering each frequency (Fourier) component of recorded signal independently. This was, for example, pursued in Miceli et al. [2017] where each frequency component of the spikes and LFP signals were computed independently (i.e., each frequency component had a specific value of *σ*_e_ and a corresponding phase shift required by the Kramers-Kronig relations to preserve causality) and eventually summed to provide the full electric potential. However, on balance the experimental evidence points to at most a weak frequency dependence of *σ*_e_ with only minor putative effects on the recorded spikes and LFPs [Miceli et al., 2017]. Therefore, the present approximation in LFPy2.0 to assume a frequency-independent conductivity *σ*_e_, seems warranted.

#### Modeling of ECoG signals

LFPy2.0 provides two different methods for computing ECoG signals, that is, signals at the cortical surface: the method-of-images (MoI) Section 2.2.2 and the four-sphere model Section 2.3.4 which both have their pros and cons. The MoI method assumes a planar cortical interface and that the media above this interface can be described electrically by means of a single isotropic electrical conductivity. The four-sphere model assumes a spherical cortical surface and uses the far-field dipole approximation which requires the dipolar sources to be sufficiently far away from the recording contacts. With the present use of current dipole moments representing entire neuron populations, this approximation is challenged by the relatively short distance between in particular the most superficial populations and the cortical surface [Næss, 2015]. A future project is to systematically explore the accuracy of these two methods for ECoG modeling, for example by comparing their predictions for different situations.

The present forward modeling of electrical potentials are based on stylized spatial (planar/spherical geometries, step-wise varying conductivities) and directional (isotropy/cylindrical anisotropy) variations. More complicated models for the variation of the extracellular conductivity can be accounted for by means of finite-element modeling (FEM, Logg et al. [2012]; Lempka and McIntyre [2013]; Ness et al. [2015]; Næss et al. [2017]) for which the ‘lead field’, that is, the contribution from transmembrane currents or dipole moments to electric signals, always can be computed [Malmivuo and Plonsey, 1995]. FEM could, for example, be used to explore in detail how the recording device affects the recorded ECoG signal when a grid of ECoG contacts are embedded in an insulating material (see, for example, Castagnola et al. [2014]), in analogy to the study of multielectrode arrays (MEAs) in Ness et al. [2015].

#### More complicated head models

The current dipole moments computed by LFPy can also be used to compute EEG and MEG signals based on geometrically detailed head models measured by MRI [Bangera et al., 2010; DeMunck et al., 2012; Vorwerk et al., 2014; Huang et al., 2016]. Note, however, that geometrically detailed head models do not automatically transfer to electrically detailed head models, and it is thus not always clear how much accuracy is gained by using such models rather than the simpler head models currently implemented in LFPy (see discussion in [Nunez and Srinivasan, 2006, Ch. 6]).

#### 4.5. Possible improvements of LFPy code

While we here demonstrated a relatively strong scaling of parallel network implementations in LFPy, the code itself could be further optimized for improving overall simulation speeds and reduced memory consumption allowing for larger networks for any given MPI pool size.

One common way of improving efficiency of Python applications is rewriting ‘slow’ code to use Cython (C-extensions for Python, http://cython.org, Smith [2015]). The current LFPy version uses Cython to a limited extent, but remaining code bottlenecks could be identified and addressed accordingly. One potential problem with efficient porting of parts of LFPy’s Python code to Cython is repeated calls to NEURON’s Python interface, which from a performance point of view should be avoided.

One known bottleneck with parallel implementations of multicompartment neuron networks is uneven load balance, resulting from the fact that individual neurons with very uneven numbers of compartments may be assigned to the different MPI processes. Uneven load balance could potentially be addressed by incorporating the multi-split method described in Hines et al. [2008], as it appears compatible with the presently used `CVode.use_fast_imem` method (available since NEURON v7.4). LFPy could then be updated accordingly.

Even without the NEURON multi-split method, distribution of cells among MPI processes using a round-robin scheme could, however, be optimized to level out large differences in compartment counts (and corresponding numbers of state variables). Memory consumption could also be addressed by choosing more efficient memory structures or generators, for example, for connectivity management, and by avoiding in-memory storage of output data wherever possible. File-based I/O operations during ongoing simulations may, however, come at the expense of increased simulation times.

In terms of improved support for simulator-independent (agnostic) model description languages for neuronal models such as NeuroML [Gleeson et al., 2010; Cannon et al., 2014] or NESTML [Plotnikov et al., 2016], LFPy’s `TemplateCell` and `NetworkCell` classes already now support loading of active and passive single-neuron model files translated to NEURON’s HOC and NMODL languages from NeuroML and NeuroML2 (now in development). A growing number of such single-neuron models is becoming available through, for example the Open Source Brain initiative (http://www.opensourcebrain.org), which can readily be used in order to construct new network models. While certainly doable, LFPy is at present not set up for automatic loading of entire neuron networks specified in NeuroML. Also, single-cell and network models specified using LFPy could, in principle, be possible to translate into NeuroML as well, which would allow for executing such models using for example NetPyne (www.neurosimlab.org/netpyne) or LEMS [Cannon et al., 2014].

### 4.6. Other measurement modalities in LFPy

The present version of LFPy only models recording of electric and magnetic brain signals. Optical recording methods are increasingly used in neurophysiology, however, and forward-modeling of such signals would be a natural extension of the present functionality. In voltage-sensitive dye imaging (VSDi), the recorded signals reflects a weighted average of the membrane potentials, and such averages can be readily computed since the membrane voltages in all neuronal compartments are computed during a network simulation simulation [Chemla and Chavane, 2010a,b]. This must then be combined with proper forward-modeling of the propagation of the light through the brain tissue [Tian et al., 2011; Abdellah et al., 2015, 2017].

Calcium imaging has become a wide-spread method for measuring neural dynamics [Grienberger and Konnerth, 2012]. With the use of neuron models that explicitly includes dynamic modelling of the intracellular calcium concentrations (for example, Hay et al. [2011]; Almog and Korngreen [2014]) such signals could be directly modeled as well.

### 4.7. Outlook

While information in the brain might largely be represented by spike trains, we believe that tools such as LFPy will be instrumental in testing candidate network models aiming to account for this information processing. In the foreseeable future, experimental data against which candidate models can be tested will be a limiting factor. It is thus key that such candidate models can be tested not only against spike trains, but also other measurement modalities.

This updated version of LFPy makes a major step towards being a true multiscale simulator of neural circuits, allowing for flexible incorporation of highly detailed neuron models at the micrometer scale, yet able to also predict recorded signals such as EEG and MEG at the systems-level scale. The largest network considered in the here had 57,600 neurons. With the present code, not optimized for numerical efficiency, the simulation of 1.5 seconds of biological time on this network required about 1600 CPU hours across 2400 MPI processes. With optimized code, we expect that much larger networks can soon be addressed routinely as ever more powerful computers gradually become available. The software is also publicly available on GitHub and retains the open-source software license of its initial release, and our hope is that continued development remains driven by needs and contributions of individuals and groups of researchers.

## 5. Acknowledgements

This work received funding from the European Union Horizon 2020 Research and Innovation Programme under Grant Agreement No. 720270 [Human Brain Project (HBP) SGA1], the Norwegian Ministry of Education and Research through the SUURPh Programme and the Norwegian Research Council (NFR) through COBRA, CINPLA and NOTUR -NN4661K. We thank Tuomo Mäki-Marttunen for useful comments and discussions on the manuscript.

